# The suboptimality of perceptual decision making with multiple alternatives

**DOI:** 10.1101/537068

**Authors:** Jiwon Yeon, Dobromir Rahnev

## Abstract

It is becoming widely appreciated that human perceptual decision making is suboptimal but the nature and origins of this suboptimality remain poorly understood. Most past research has employed tasks with two stimulus categories, but such designs cannot fully capture the limitations inherent in naturalistic perceptual decisions where choices are rarely between only two alternatives. We conducted four experiments with tasks involving multiple alternatives and used computational modeling to determine the decision-level representation on which the perceptual decisions were based. The results from all four experiments pointed to the existence of robust suboptimality such that most of the information in the sensory representation was lost during the transformation to a decision-level representation. These results reveal severe limits in the quality of decision-level representations for multiple alternatives and have strong implications about perceptual decision making in naturalistic settings.

## Introduction

Perception has been conceptualized as a process of inference for over a century and a half^1^. According to this view, the outside world is encoded in a pattern of neural firing and the brain needs to decide what these patterns signify. Hundreds of papers have revealed that this inference process is suboptimal in a number of different ways^2^. However, these papers have almost exclusively employed tasks with only two stimulus categories (though notable exceptions exist^3,4^). Experimental designs where decisions are always between two alternatives cannot fully capture the processes inherent in naturalistic perceptual decisions where stimuli can belong to many different categories (e.g., which of all possible local species does a particular tree belong to). Therefore, fully understanding the mechanisms and limitations of perceptual decision making requires that we characterize the process of making decisions with multiple alternatives.

One critical difference between perceptual decisions with two versus multiple alternatives is the richness of the sensory information that the decisions are based on. Decisions with two alternatives can be based on the evidence for each of the two categories, or even just the difference between these two pieces of evidence^5^. For example, traditional theories such as signal detection theory^6^ and the drift diffusion model^7^ postulate that 2-choice tasks are performed by first summarizing the evidence down to a single number – the location on the evidence axis in signal detection theory and the identity of the boundary that is crossed in drift diffusion – that is subsequently used for decision making. Thus, the sensory information relevant for the decision in such tasks is relatively simple and could potentially be represented in decision-making circuits without substantial loss of information. However, the relevant sensory information in multi-alternative decisions is more complex because it contains the evidence for each of the multiple alternatives available. Further, this richer sensory information can no longer be summarized in a simple form in decision-making circuits without a substantial loss of information (**Figure 1**). However, it is currently unknown whether decision-making circuits can represent the rich information from sensory circuits in the context of multi-alternative decisions or whether the decision-making circuits only represent a crude summary of the sensory representation.

**Figure 1.**
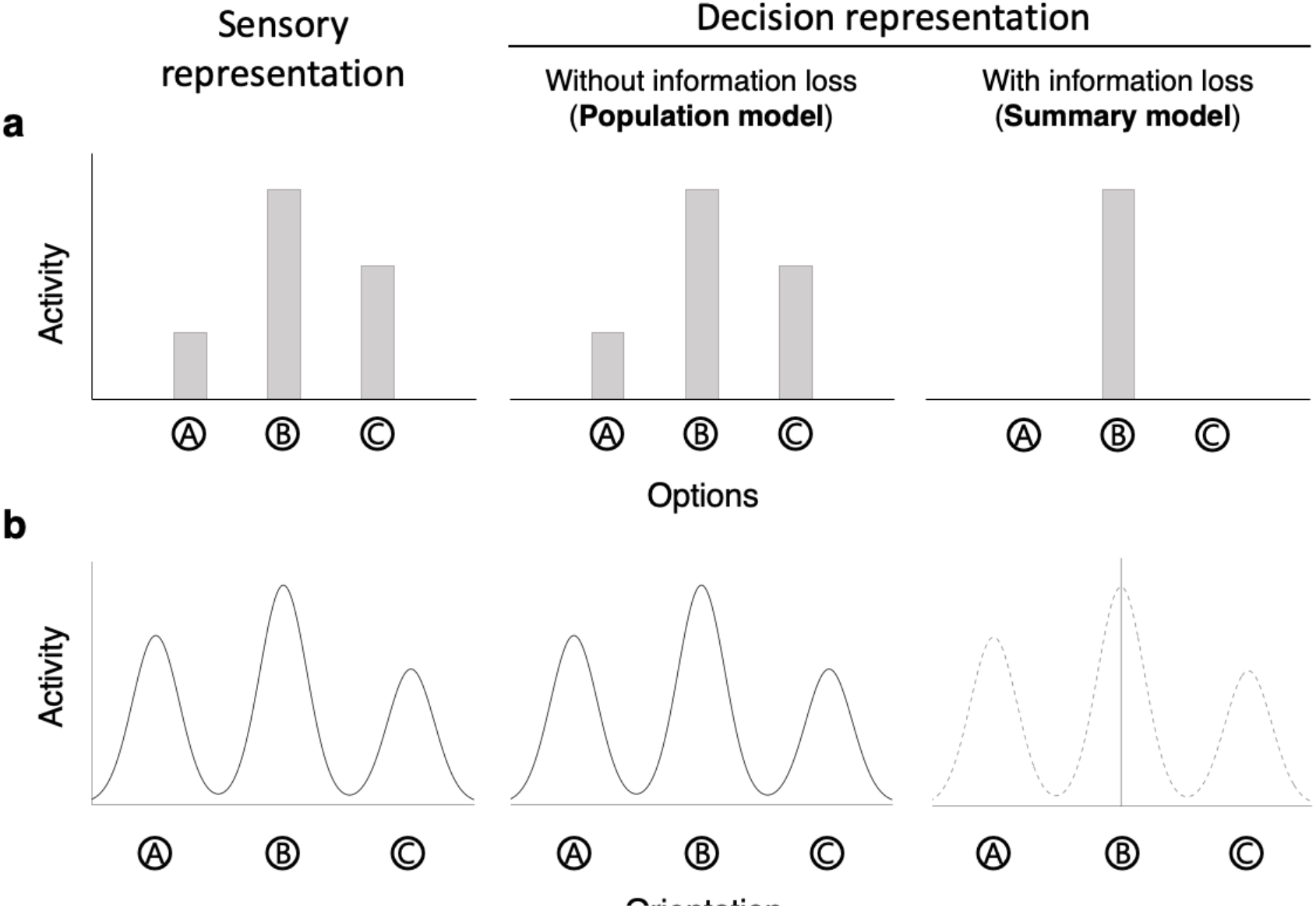
Sensory and decision-level representations in perceptual decision making with multiple alternatives. (a) Decision making with multiple discrete alternatives. In cases where a subject has to choose between multiple discrete alternatives (e.g., options A, B, and C), a stimulus can be assumed to give rise to a sensory representation that consists of different amount of sensory activity for each alternative (left panel). A decision-level representation without information loss would consist of a copy for the sensory representation (middle panel). We refer to this possibility as a ‘population’ model of decision-level representation. On the other hand, a decision-level representation may consist of only a summary of the sensory representation thus incurring information loss. One possible summary representation consists of passing only the highest activity onto decision-making circuits (right panel). We refer to this type of representation as a ‘summary’ model of decision-level representation. This summary representation involves information loss that will become apparent if subjects have to choose between the other alternatives (e.g., alternatives A and C). (b) Decision making with continuous but multimodal sensory representation. Similar to having multiple discrete alternatives, decisions can involve judging a continuous feature (e.g., orientation) but in the context of a multimodal (e.g., a trimodal) underlying sensory representation (left panel). The decision-level representation can again consist of either a copy for the sensory representation (middle panel) or a summary of this sensory representation (right panel).

To uncover the decision-level representation of decisions with multiple alternatives, we used discrete stimulus categories in three experiments and stimuli that give rise to a trimodal sensory distribution in a fourth experiment. All experiments featured a condition where subjects picked the dominant stimulus among all of the possible stimulus categories (four different colors in Experiment 1, six different symbols in Experiments 2 and 3, and three different stimulus direction in Experiment 4). Based on these responses, we estimated the parameters of a model describing subjects’ internal distribution of sensory responses (that is, the activity levels for each stimulus category). We then included conditions where subjects were told to pick between only two alternatives after the offset of the stimulus (Experiments 1, 2, and 4) or to make a second choice if the first one was incorrect (Experiment 3). These conditions allowed us to compare different models of how the sensory representation was transformed into a decision-level representation. To anticipate, we found robust evidence for suboptimality in that decisions in our experiments were based on a summary of the sensory representation thus incurring substantial information loss. These results indicate that perceptual decision-making circuits may not have access to the full sensory representation in the context of multiple alternatives and that significant amount of simplification is likely to occur before sensory information is used for deliberate decisions.

## Results

We investigated whether the decision-level representation in perceptual decision making with multiple alternatives consists of the whole sensory representation or only a summary of it. To address this question, we performed four experiments in which subjects made choices about discrete stimulus categories or continuous variables giving rise to multimodal sensory distributions.

### Experiment 1

Experiment 1 required subjects to pick which of four possible colors – blue, red, green, and white – was most frequently presented (**Figure 2**). The stimulus consisted of 49 colored circles arranged in a 7×7 square presented for 500 ms. On each trial, one color was randomly chosen to be “dominant” and 16 circles were painted in that color, whereas the remaining three colors were “non-dominant” and 11 circles were painted in each of those colors. The experiment featured two different conditions. In the 4-alternative condition, subjects picked the dominant color among the four possible colors. In the 2-alternative condition, after the offset of the stimulus, subjects were asked to choose between the dominant and one randomly chosen non-dominant color. In both conditions, the response screen was displayed with 0-ms delay thus minimizing short-term memory demands. Note that subjects’ task was always to correctly identify the dominant color.

**Figure 2.**
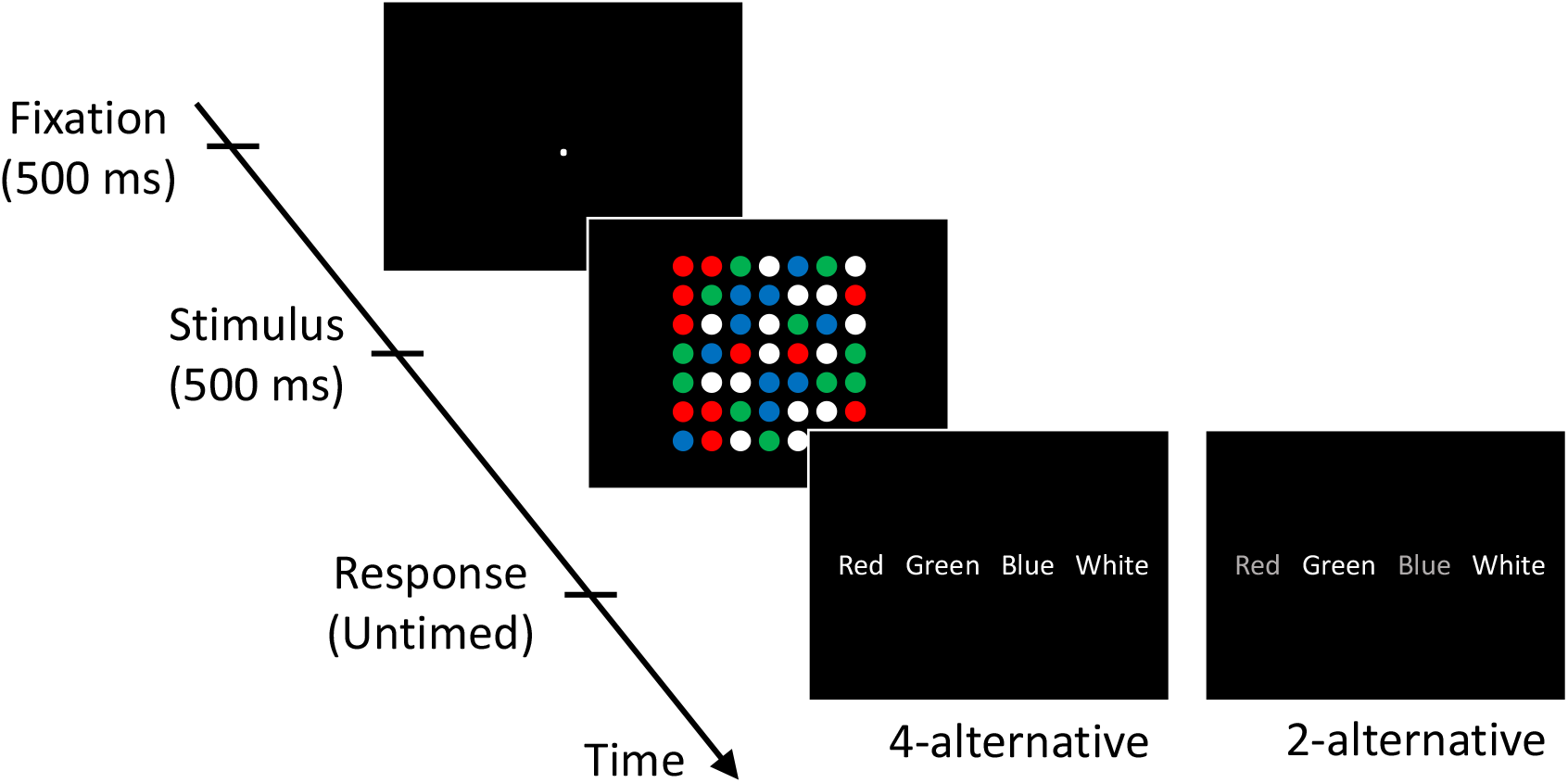
Task for Experiment 1. Each trial consisted of a fixation period (500 ms), stimulus presentation (500 ms), and untimed response period. The stimulus comprised of four different colored circles (red, green, blue, and white). One of the colors (white in this example) was presented more frequently (16 circles; dominant color) than the other colors (11 circles each; non-dominant colors). Subjects’ task was to indicate the dominant color. Two conditions were presented in different blocks. In the 4-alternative condition, subjects chose between all four colors. In a separate 2-alternative condition, on each trial subjects were given a choice between the dominant and one randomly chosen non-dominant color.

Using subjects’ responses in the 4-alternative condition, we estimated the parameters of the sensory distribution representing the activity level for each color. We then considered the predictions for the 2-alternative condition of two different models: (1) a ‘population’ model, according to which perceptual decisions are based on the whole distribution of activities over the four colors, and (2) a ‘summary’ model, according to which perceptual decisions are based on a summary of the whole distribution. There are a number of ways to create a summary of the whole distribution. However, in the context of this task, the only relevant information is the order of activation levels from highest to lowest (this order determines how a subject would pick different colors as the dominant color in the 2-alternative condition). Other information, such as average activity level, is irrelevant to the task here. We first considered an extreme summary model that consists of the activity level for the one color with highest level of activity. Other summary models, in which decision-making circuits have access to the activity levels of the n>1 colors with highest activity levels, are examined later.

The population and summary models could be easily compared because they make different predictions about a subject’s performance in the 2-alternative task (for a mathematical derivation, see Supplementary Methods). Indeed, the models make the same prediction when the dominant color gives rise to the highest activity level (**Figure 3a**) and when the alternative option given to the subject happens to have the highest activity (**Figure 3b**), but diverge when the highest activity is associated with a color that is not among the two options with the population model predicting a higher performance level (**Figure 3c**).

**Figure 3.**
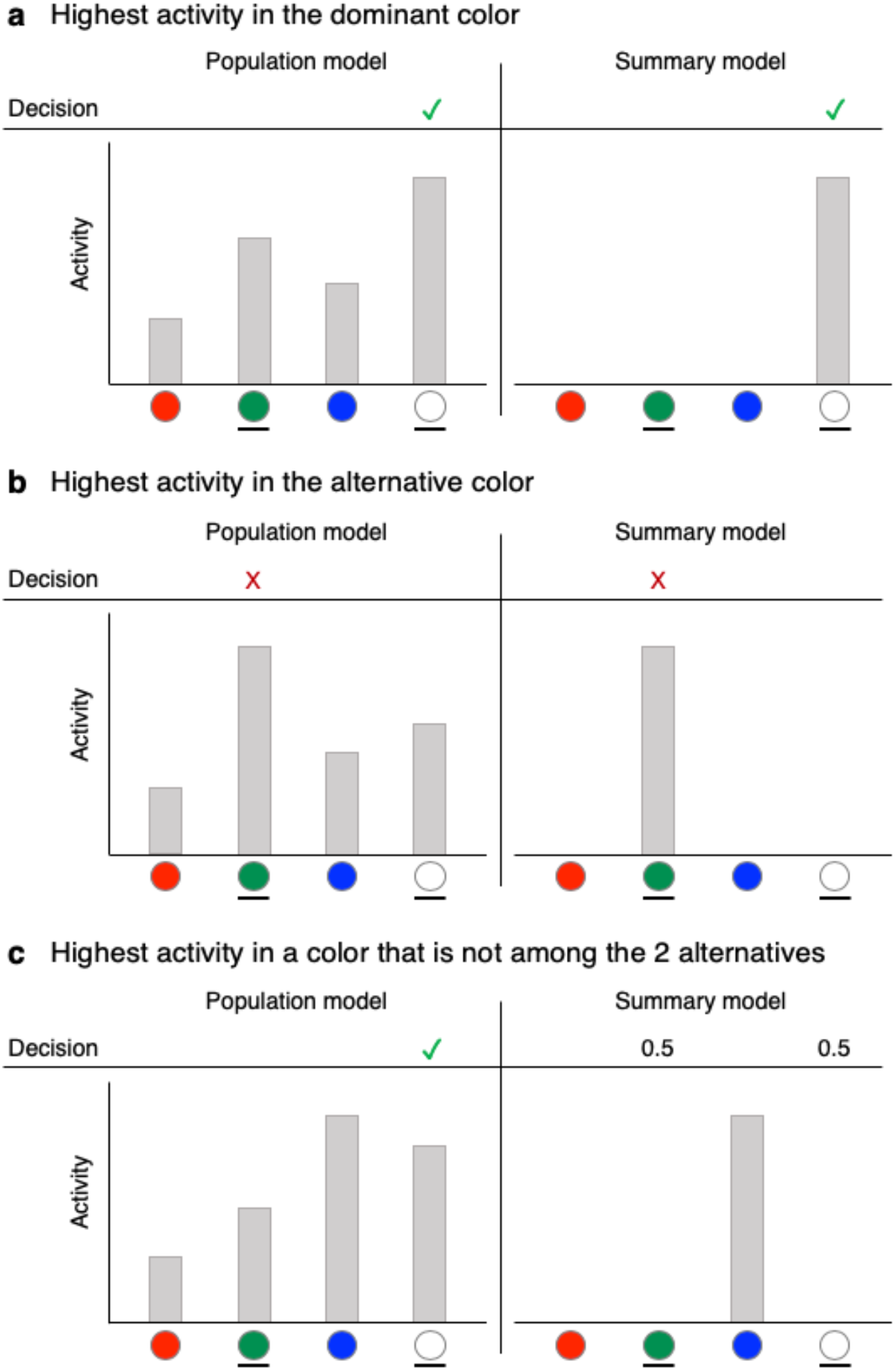
Predictions of the population and summary models for subjects’ choices in the 2-alternative condition. The population model (left panels) assumes that decision-making circuits have access to the activity levels associated with each of the four colors (four gray bars), whereas the summary model (right panels) assumes that decision-making circuits only have access to the highest activity level (single gray bar). In all examples, the dominant circle is white, and subjects are given a choice between white and green. (a) When the highest activity happens to be at the dominant color, both models predict that the subject would correctly choose the dominant color. (b) When the highest activity happens to be at the alternative color, both models predict that the subject would incorrectly choose the alternative color. (c) The two models’ prediction diverge when the highest activity is associated with a color other than the two presented alternatives. In such cases, the activation for the dominant color is likely to be higher than for the alternative color, so according to the population model, subjects would ignore the color with the highest activity (red color in the example here) and correctly pick the dominant color in the majority of the trials. However, according to the summary model, subjects have no information about the activation levels for the dominant and the alternative colors and would thus correctly pick the dominant color on only 50% of such trials.

The difference between the two models could be seen in the actual model predictions. Indeed, based on the performance in the 4-alternative condition (average accuracy = 69.2%, chance level = 25%), the population and summary models predicted an average accuracy of 84.2% and 79.7% in the 2-alternative condition, respectively. Compared to the actual subject performance (average accuracy = 78%), the population model overestimated the accuracy in the 2-alternative conditions for 29 of the 32 subjects (average difference = 6.21%, t(31) = 8.19, *p* = 3.02 x 10^−9^). Surprisingly, the summary model also overestimated the accuracy in the 2-alternative condition but the misprediction was much smaller (average difference = 1.72%, t(31) = 2.35, *p* = .025) (**Figure 4a**). Indeed, the absolute error of the predictions of the population model (average = 6.54%) was significantly larger than for the summary model (average = 3.61%; t(31) = 5.65, *p* = 3.34x 10^−6^). Overall, the summary model predicted the accuracy in the 2-alternative condition better than the population model for 26 of the 32 subjects (**Figure 4b**).

**Figure 4.**
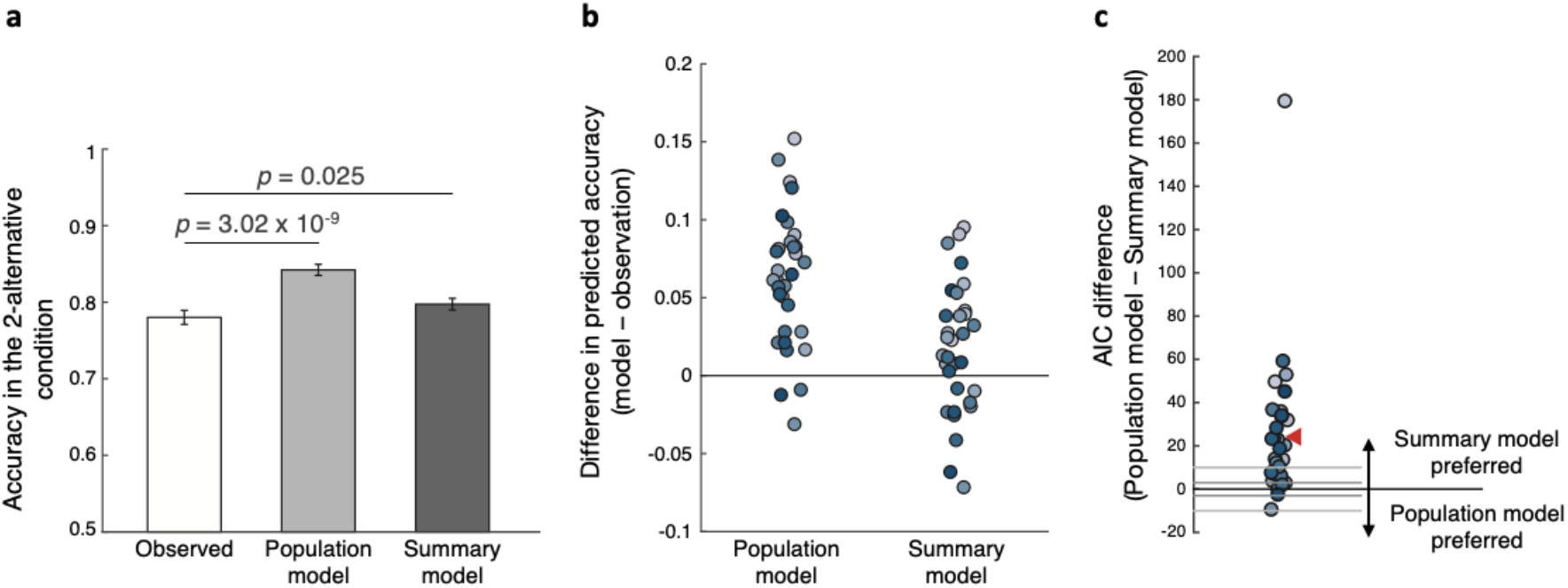
Comparisons between the population and summary models in Experiment 1. (a) Task accuracy in the 2-alternative condition observed in the actual data (white bar), and predicted by the population (light gray bar) and summary (dark gray bar) models. The predictions for both models were derived based on the data in the 4-alternative condition. (b) Individual subjects’ differences in the accuracy in the 2-alternative condition between the two models and the observed data. (c) Difference in Akaike Information Criterion (AIC) between the population and the summary models. Positive AIC values indicate that the summary model provides a better fit to the data. Each dot represents one subject. The gray horizontal lines at ±3 and ±10 indicate common thresholds for suggestive and strong evidence for one model over another. The red triangle indicates the average AIC difference. The summary model provided a better fit than the population model for 30 of the 32 subjects.

We further compared the models’ fits to the whole distribution of responses. We found that the Akaike Information Criterion (AIC) favored the summary model by on average 24.30 points (**Figure 4c**), which corresponds to the summary model being 1.89 x 10^5^ times more likely than the population model for the average subject. Across the whole group of 32 subjects, the total AIC difference was thus 777.63 points, corresponding to the summary model being 7.26 x 10^168^ times more likely in the group. Note that since the population and summary models had the same number of parameters, the same results would be obtained regardless of the exact metric employed (e.g., the BIC differences would be exactly the same).

Finally, we constructed and tested four additional models. The first two models postulated that decision-making circuits have access to the two or three highest activations of the sensory distribution (“2-highest” and “3-highest” models, respectively). These models could thus be seen as intermediate options between the summary and population models. Two other models postulated that subjects choose either two or three stimulus categories to attend to and then make their decisions based on a full probability distribution over the activity levels of the attended categories (“2-attention” and “3-attention” models; see **Supplementary Methods** for details). We found that all of these models produced poor fits to the 2-alternative condition and were outperformed by the summary model (**Supplementary Results** and **Supplementary Figures 1, 6a-d**).

### Experiment 2

The results from Experiment 1 strongly suggest that within the context of our experiment, decision-making circuits do not represent the whole sensory distribution but only a summary of it. We sought to confirm and generalize these findings in two additional, pre-registered experiments. For Experiment 2, we made several modifications: (1) we changed the stimulus from color to symbols, (2) we raised the number of stimulus categories from four to six, and (3) we significantly increased the number of trials per subject in order to obtain stronger results on the individual-subject level. Specifically, we presented the six symbols ‘?’, ‘#’, ‘$’, ‘%’, ‘+’, and ‘>’ such that the dominant symbol was presented 14 times and each non-dominant symbol was presented 7 times (**Figure 5a**). The 49 total symbols were again arranged in a ×7 grid. Each subject completed a total of 3,000 trials that included equal number of trials of a 6-alternative condition and a 2-alternative condition that were equivalent to the 4- and 2-alternative conditions in Experiment 1.

**Figure 5.**
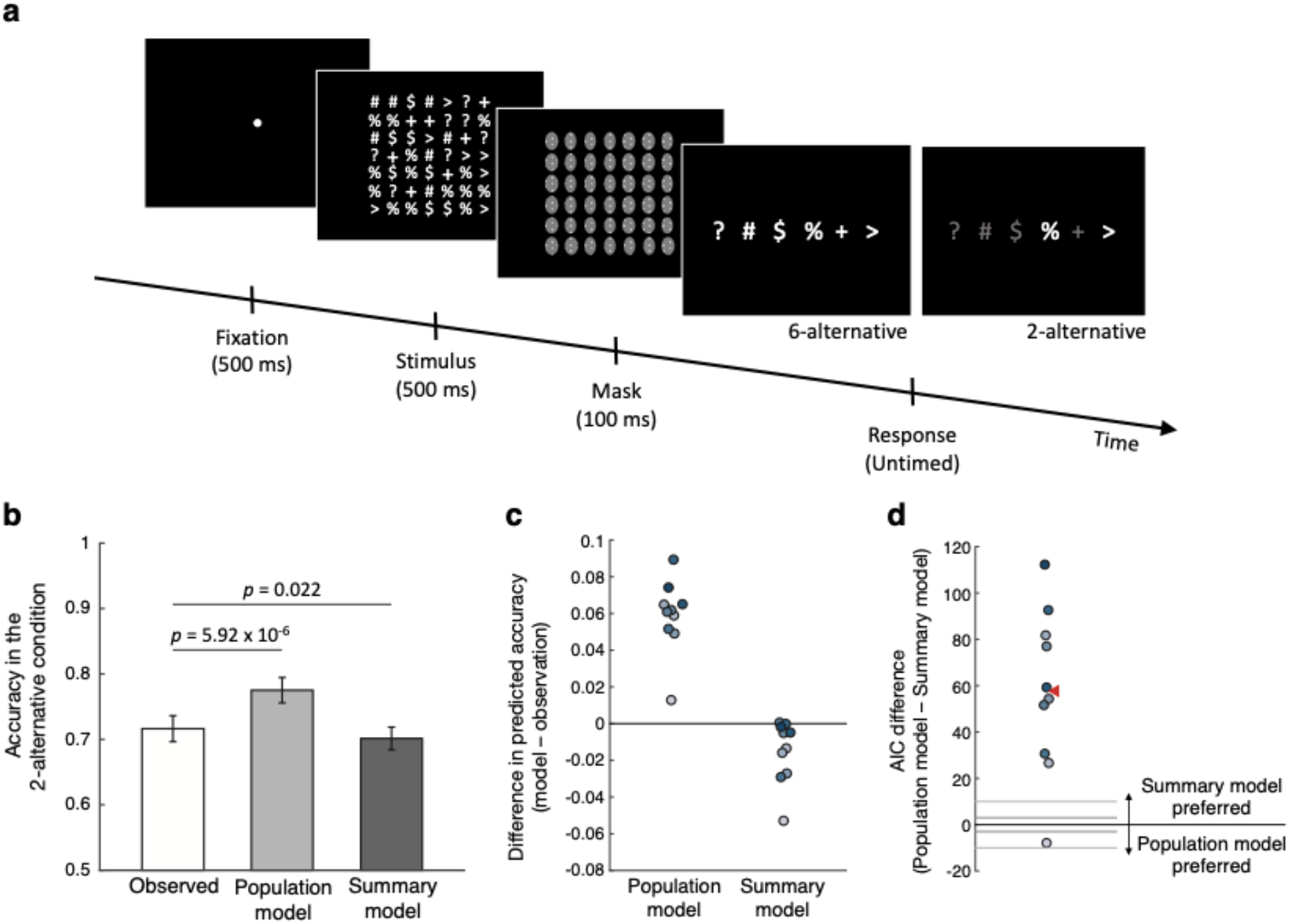
Task and results for Experiment 2. (a) The task in Experiment 2 was similar to Experiment 1 except for using six different symbols (‘?’, ‘#’, ‘$’, ‘%’, ‘+’, and ‘>’) instead of four different colors. One of the symbols was presented more frequently (14 times, dominant symbol) than the others (7 times each, non-dominant symbols) and subjects’ task was to indicate the dominant symbol. Two conditions were presented in different blocks: a 6-alternative condition where subject chose between all six symbols and a 2-alternative condition where subjects were given a choice between the dominant and one randomly chosen non-dominant symbol. (b) Task accuracy in the 2-alternative condition observed in the actual data (white bar) and predicted by the population (light gray bar) and summary (dark gray bar) models. The predictions for both models were derived based on the data in the 6-alternative condition. (c) Individual subjects’ differences in the accuracy of the 2-alternative condition between the two models and the observed data. (d) Difference in Akaike Information Criterion (AIC) between the population and the summary models. Positive AIC values indicate that the summary model provides a better fit to the data. Each dot represents one subject. The red triangle indicates the average AIC difference. The summary model provided a better fit than the population model for nine out of 10 subjects.

Just as in Experiment 1, we computed the parameters of the sensory representation using the trials from the 6-alternative condition (average accuracy = 50.5%, chance level = 16.7%) and used these parameters to compare the population and summary models’ predictions for the 2-alternative condition. We found that the average accuracy in the 2-alternative condition (71.6%) was slightly underestimated by the summary model (predicted accuracy = 70.1%, t(9) = 2.76, *p* = .022) but was again significantly overestimated by the population model (predicted accuracy = 77.5%, t(9) = 9.41, *p* = 5.92 x 10^−6^) (**Figure 5b**). Individually, the summary model provided better prediction of the accuracy in the 2-alternative condition for nine out of the 10 subjects (**Figure 5c**).

Further, we compared the population and summary models’ fits to the whole distribution of responses. We found that the summary model was preferred nine of our 10 subjects and the difference in AIC values in all these nine subjects were larger than 25 points (**Figure 5d**). The AIC values of the one subject for whom the population model was favored over the summary model differed only by 7.9 points. On average, the summary model had an AIC value that was 57.79 points lower than the population model corresponding to the summary model being 3.55 x 10^12^ times more likely for the average subject. Across the whole group of 10 subjects, the total AIC difference was thus 577.94 points, corresponding to the summary model being 3.14 x 10^125^ times more likely in the group. Finally, we found that the additional four models again mispredicted the performance in the 2-alternative condition and were outperformed by the summary model (**Supplementary Results** and **Supplementary Figures 2, 6e-h**).

### Experiment 3

Taken together, Experiments 1 and 2 suggest that in the context of multi-alternative decisions, the system for deliberate decision making may not have access to the whole sensory representation. This conclusion is based on experiments that differed in the nature of the stimulus, the number of stimulus categories, and the amount of trials that subjects performed. Nevertheless, both Experiments 1 and 2 relied on the same design of comparing 4- (or 6-) and 2-alternative conditions. Therefore, to further establish the generality of our results, in Experiment 3 we employed a different experimental design. We used the same stimulus as in Experiment 2 and presented all 6 alternatives on every trial, but additionally gave subjects the opportunity to provide a second answer on about 40% of error trials (**Figure 6a**). Using the performance on the first answer, we compared the predictions of the population and summary models for the second answers.

**Figure 6.**
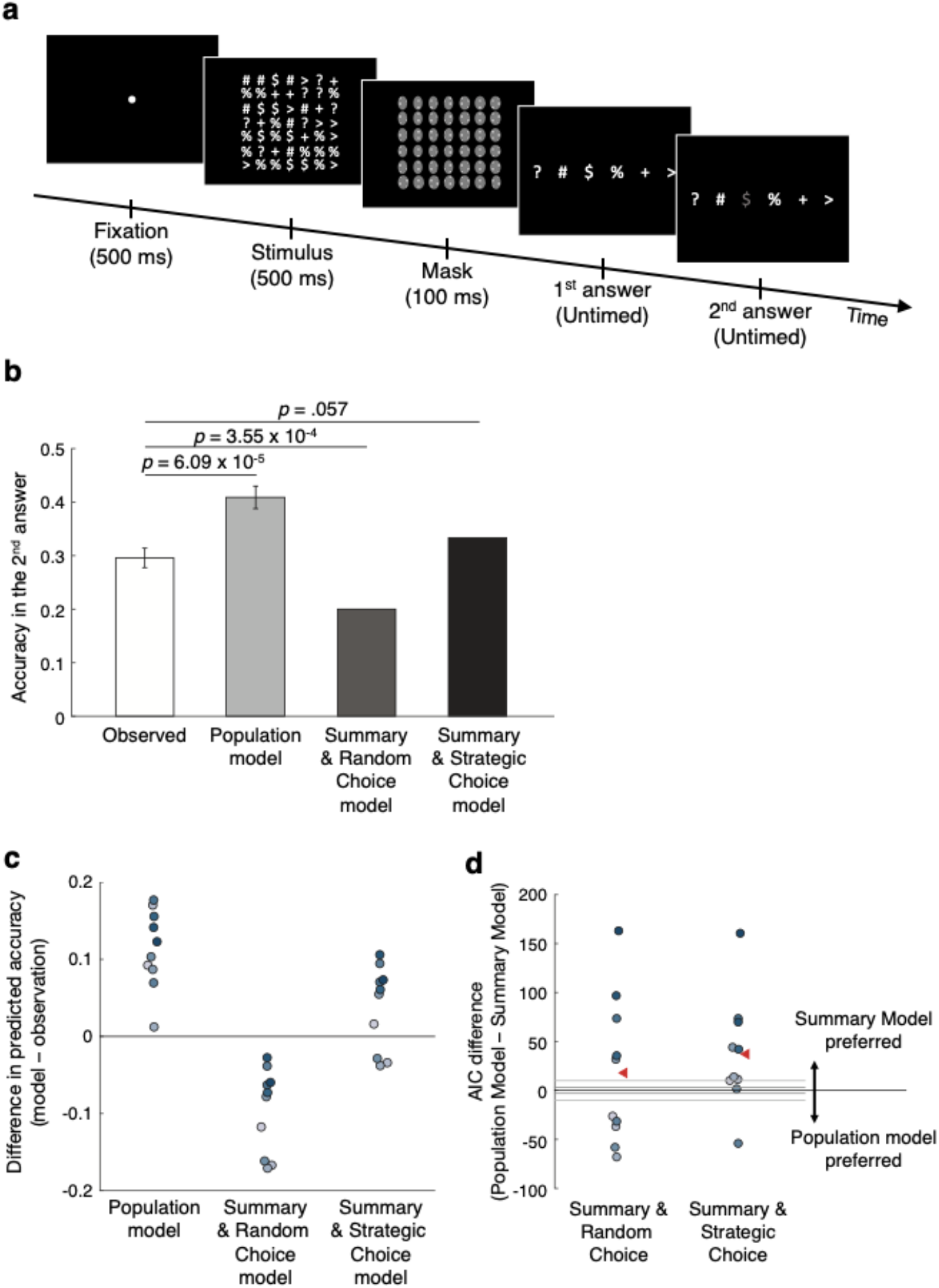
Task and results for Experiment 3. (a) The same stimuli as in Experiment 2 were used in Experiment 3 but the task was slightly different. Subjects always reported the dominant symbol among all six alternatives. However, on 40% of the trials in which they gave a wrong answer, subjects were given the opportunity to make a second guess. (b) Task accuracy for the second answer observed in the actual data (white bar), predicted by the population model (light gray bar), predicted by the Summary & Random Choice model (dark gray bar), and predicted by the Summary & Strategic Choice model (black bar). The predictions of the three models were derived based on subjects’ first answers. (c) Individual subjects’ differences in the accuracy of the second answer between each model’s prediction and the observed data. (d) Difference in Akaike Information Criterion (AIC) between the population and the two summary models. Positive AIC values indicate that the summary model provides a better fit to the data. Each dot represents one subject. The red triangle indicates the average AIC difference.

The population model makes a clear prediction about the second answer – subjects should choose the stimulus category with the highest activation from among the remaining five options. The second answer will thus have relatively high accuracy because the presented stimulus category is likely to produce one of the highest activity levels (**Supplementary Figure 3a**). Note that in the context of this experiment, the 2-Highest and 3-Highest models are functionally equivalent to the population model since they both represent the category with the second highest activity, and both allow that this stimulus category be chosen with the second answer.

On the other hand, the summary model only features information about the stimulus category with the highest activity. Once that stimulus category is chosen as the first answer, the model postulates that the subject does not have access to the activations associated with the other stimulus categories. Given this representation, subjects could adopt at least two different response strategies. One possible strategy is for the subject to make their second answer at random, which would result in chance level (20%) performance. We call this the “Summary & Random Choice” model (**Supplementary Figure 3b**). However, another possibility is for the subject to make the second answer strategically. One available strategy is for the subject to pick the stimulus category of a randomly recalled symbol from the ×7 grid. Given that subjects inspected the stimuli for 500 ms, they could easily remember one location with a symbol other than the one they picked for their first answer. We call this the “Summary & Strategic Choice” model (**Supplementary Figure 3c**). According to this model, the second answer will be correct on 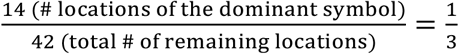 or 33.3% of the time. Conversely, each of the four remaining incorrect categories will be chosen on 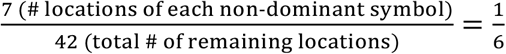 or 16.7% of the time (**Supplementary Figure 4**).

To adjudicate between these three models, we first examined subjects’ accuracy on the first answer. Subjects responded correctly in their first answer on 50.7% of the trials (chance level = 16.7%). Using this performance, we computed the parameters of the sensory representation as in Experiments 1 and 2 in order to generate the models’ predictions for the second answer. We found that subjects’ accuracy for the second answers was 29.6%. This value was greatly overestimated by the population model, which predicted accuracy of 40.9% (t(9) = 7.04, *p* = 6.09 x 10^−5^; **Figure 6b**). On the other hand, the Summary & Random Choice model greatly underestimated the observed accuracy (predicted accuracy = 20%, t(9) = 5.55, *p* = 3.55 x 10^−4^). Finally, the Summary & Strategic Choice model produced the most accurate prediction (predicted accuracy = 33.3%, t(9) = 2.18, *p* = .057). On an individual subject level, the population model overestimated the accuracy of the second answer for all 10 subjects, the Summary & Random Choice model underestimated the accuracy of the second answer for all of the 10 subjects, whereas the Summary & Strategic Choice model was best calibrated overestimating the accuracy of the second answer for 7 subjects and underestimating it for the remaining 3 subjects (**Figure 6C**).

Formal comparisons of the models’ ability to fit the full distribution of responses for the second answers demonstrated that the population model provided the worst overall fit (**Figure 6d**). Indeed, the population model resulted in AIC values that were higher than the Summary & Random Choice model by an average of 18.05 points (corresponding to 8.29 x 10^3^-fold difference in likelihood in the average subject) and a total of 180.46 points (corresponding to 1.53 x 10^39^-fold difference in likelihood in the group). The population model underperformed the Summary & Strategic Choice model even more severely (average AIC difference = 37.29 points, corresponding to 1.25 x 10^8^-fold difference in likelihood in the average subject; total AIC difference = 372.93 points, corresponding to 9.57 x 10^80^-fold difference in likelihood in the group). Lastly, the Summary & Strategic Choice model also outperformed all of the additional models (**Supplementary Results** and **Supplementary Figure 6i-l**). Thus, just as Experiments 1 and 2, Experiment 3 provides strong evidence that in the context of multi-alternative decisions, decision-making circuits only contain a summary of the sensory representation.

### Experiment 4

To adjudicate between the population and summary models, Experiments 1-3 employed discrete stimulus categories (**Figure 1a**). However, it remains possible that the results do not generalize to stimuli represented on a continuous scale. To address this issue, we performed a fourth experiment that employed a feature (dot motion) represented on a continuous scale (degree orientation). We adapted the design of Treue, Hol and Rauber^8^ in which groups of dots slid transparently across one another. Specifically, we presented moving dot stimuli where three sets of dots moved in three different directions. As Treue et al. noted, these stimuli produce the subjective experience of seeing three distinct surfaces sliding across each other and therefore give rise to a trimodal internal sensory distribution of motion direction. In each trial, one of the motion directions was represented by more dots (“dominant” direction) than the other two (“non-dominant” directions). Subjects had to indicate the dominant direction of motion, which corresponds to the location of the tallest peak in the trimodal sensory distribution (**Figure 1b**).

We used a similar modeling approach as in the previous three experiments (see Methods) where we fitted a model of the sensory representation to the 3-alternative condition (average accuracy = 77.4%, chance level = 33.3%), and compared the predicted accuracy of the 2-alternative condition between the population and summary models. As in the previous experiments, we observed that the population model consistently overestimated the accuracy of the 2-alternative condition (observed accuracy = 83.7%; predicted accuracy = 85.9%, t(10) = 4.31, *p* = .002), whereas the summary model predicted the observed accuracy well (predicted accuracy = 83.1%, t(10) = 1.37, *p* = .2) (**Figure 7b**). In a direct comparison between the two models, the summary model predicted the observed task accuracy better for nine out of 11 subjects (**Figure 7c**).

**Figure 7.**
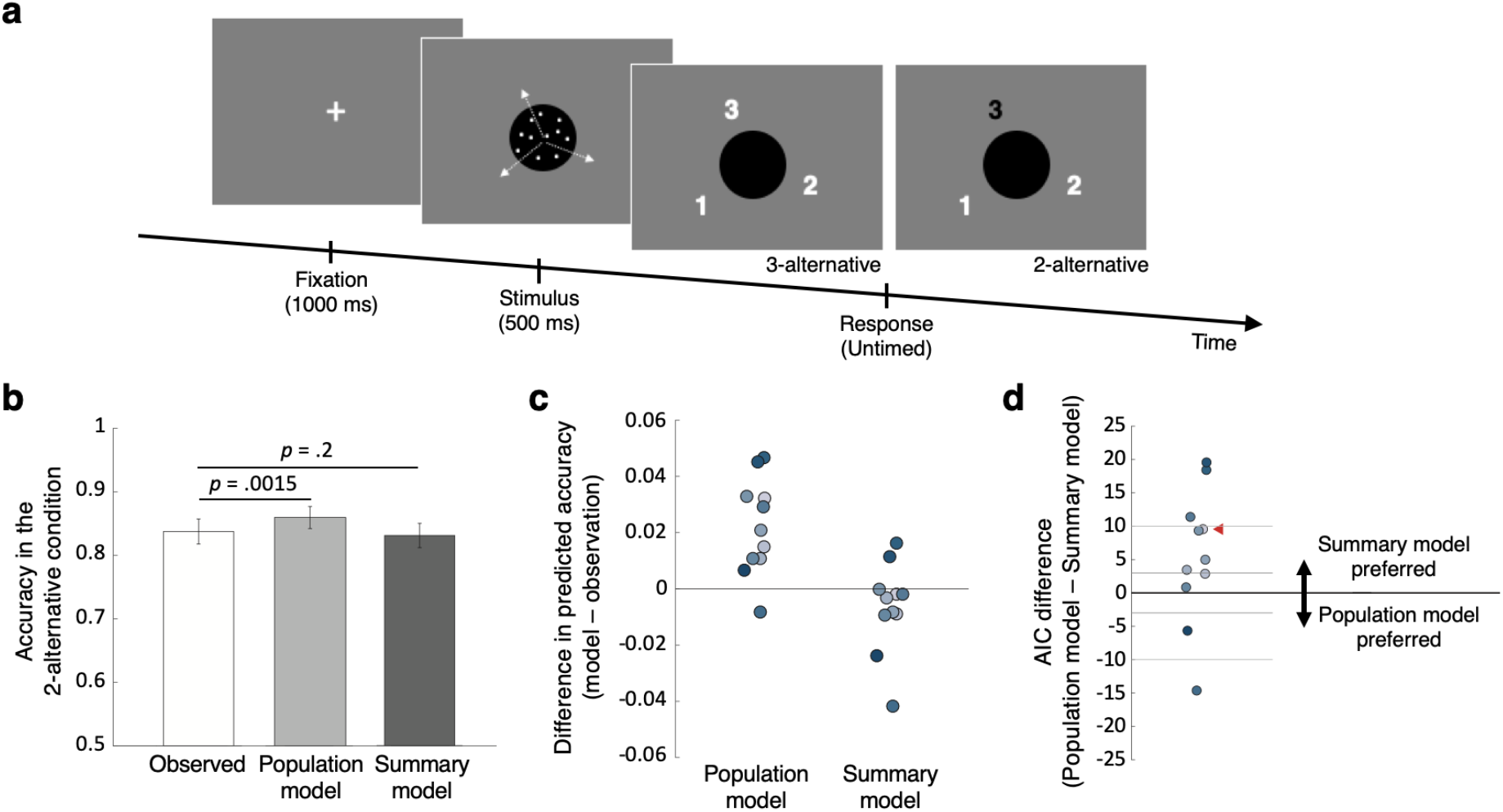
Task and results for Experiment 4 which uses stimuli (moving dots) represented on a continuous scale. (a) Three sets of dots moved in three different direction separated by 120°. Similar to Experiments 1-3, one of the three sets of dots had more dots (“dominant” direction) compared to the other two sets (“non-dominant” directions). Each trial began with a fixation cross followed by the moving dot stimulus presented for 500 ms. The response screen was presented immediately after the offset of the moving dots and randomly assigned a stimulus-response mapping on each trial. Similar to Experiments 1 and 2, subjects picked the dominant direction of motion between all three directions (3-alternative condition) or between the dominant direction and one randomly chosen non-dominant direction (2-alternative condition). (b) Task accuracy in the 2-alternatvie condition observed in the actual data (white bar) and predicted by the population (light gray bar) and summary (dark gray bar) models. The predictions for both models were derived based on the data in the 3-alternative condition. (c) Individual subjects’ differences in the accuracy of the 2-alternative condition between the predictions of each of the two models and the observed data. (d) Difference in Akaike Information Criterion (AIC) between the population and summary models. Positive AIC values indicate that the summary model predicts the observed data better. The red triangle indicates the average AIC difference. The summary model fits better than the population model for nine out of the 11 subjects.

Finally, we compared the population and summary models’ fits to the whole distribution of responses. On average, the AIC value of the summary model was lower by 5.47 points than the population model corresponding to the summary model being 15.40 times more likely for the average subject (**Figure 7d**). The total AIC difference across all subjects was 60.15 points lower for the summary model, corresponding to the summary model being 1.15 x 10^13^ times more likely than the population model.

## Discussion

We investigated whether the decision-level representation in decisions with multiple alternatives consists of a copy of the sensory representation or only a summary of it. We performed four experiments with either discrete stimulus categories or continuous stimuli producing multimodal distributions. The results across all experiments showed that the population model that assumes no loss of information from sensory to decision-making circuits did not provide a good fit to the data. Instead, the summary model, which assumes that decision-making circuits represent a reduced form of the sensory distribution, consistently provided a substantially better fit. These results strongly suggest that deliberate decision making for multiple alternatives only has access to a summary form of the sensory representation.

Prior studies have convincingly demonstrated that humans can form complex, non-Gaussian^9^ and even bimodal^10^ priors over repeated exposures to a given stimulus. However, it should be emphasized that this previous research has focused on the ability to learn a prior over many trials and did not examine the ability to use the sensory representation produced by a single stimulus on a single trial. To the best of our knowledge, the current experiments are the first to address the question of whether complex sensory codes for a single stimulus can be accurately represented in decision-making circuits.

Why is it that decision-making circuits do not maintain a copy of the full sensory representation? While our study does not directly address the origins of this suboptimality, it is likely that the reason for it lies in decision-making circuits having markedly smaller bandwidth than sensory circuits. One potential reason for this is that decision-making circuits have to be able to represent many different features (e.g., orientation, color, shape, object identity, etc.) each of which is processed in dedicated sensory regions. According to this line of reasoning, the generality inherent in decision-level representations necessitates that detail present in sensory cortex is lost. A related reason for the information loss in decision-making circuits is that the representations in such circuits often need to be maintained over a few seconds and therefore are subject to the well-known short-term memory decay^11–14^. For example, even though all conditions in our task were shown with a 0-ms delay, information may need to be passed serially from sensory to decision-making circuits thus inherently inducing short-term memory demands. According to this line of reasoning, even if decision-making circuits can represent a full copy of the information in sensory cortex, that copy will decay as soon as it begins to be assembled and thus information loss with necessarily accrue before the representation can be used for subsequent computations. Therefore, both the necessary generality of decision-level representations and the inherent limitations of short-term memory likely contribute to the sparse representations in decision-making circuits.

If decision-making circuits indeed only have access to a summary form of the sensory representation, does that mean that absolutely no computations can be based on the complete sensory representations? There is evidence that decisions can take into account the full sensory representation both in simple situations where only two alternatives are present and in cases of automatic multisensory integration^3,15–19^. In general, automated computations performed directly on the sensory representations may use the whole sensory representation^19^. Therefore, certain types of decision making can be performed via mechanisms that do indeed take advantage of the entire sensory representation with either no or minimal loss of information. However, it is likely that such computations are restricted to either very simple decisions or processes that are already automated. On the other hand, the last stage of decision-making supporting non-automatic, flexible, and deliberate decisions (which require short-term memory maintenance) only has access to a summary of the sensory representation.

Our findings can be misinterpreted as suggesting that complex visual displays are represented as a point estimate: that is, the decision-level representation features only the best guess of the system (e.g., “60°” orientation, “red” color, or “+” symbol). The possibility of a decision-level representation consisting of a single point estimate has been thoroughly debunked^20–24^. For example, a point estimate does not allow us to rate how confident we are in our decision because we lack a sense of how uncertain our point estimate is. Given that humans and animals can use confidence ratings to judge the likely accuracy of their decisions^25–27^, decision-making circuits must have access to more than a point estimate of the stimulus.

It should therefore be clarified that our summary model does not imply that decision making operates on point estimates. Indeed, as conceptualized in Figure 1, the summary model assumes that subjects have access to both the identity of the most likely stimulus category (e.g., the color “white”) and the level of activity associated with that stimulus category. The level of activity can then be used as a measure of uncertainty, and confidence levels can be based on this level. Such confidence ratings will be less informative than the perceptual decision, which is exactly what has been observed in a number of studies^2,28,29^. In addition, this type of confidence generation may explain findings that confidence tends to be biased towards the level of the evidence for the chosen stimulus category and tends to ignore the level of evidence against the chosen category^30–35^. Thus, a summary model, consisting of the identity of the most likely stimulus and the level of activity associated with this stimulus, appears to be broadly consistent with findings related to how people compute uncertainty and is qualitatively different than a decision-level representation consisting of a point estimate.

Another important question concerns whether any additional information is extracted from the sensory representation beyond what is assumed by the summary model. It is well known that humans can quickly and accurately extract a high-order “gist” of a scene^36–38^, as well as the statistical structure of an image^39^. Therefore, it appears that rich information is extracted during the time when the stimulus is being viewed. In fact, this information often goes beyond the extraction of just the identity of the most likely stimulus and the level of activity associated with this stimulus assumed by our summary model. For example, our moving dots stimulus in Experiment 4 resulted in the perception of three surfaces sliding on top of each other. This means that what was extracted in that experiment was the approximate location of each of the three peaks of the trimodal sensory distribution. Similarly, the subjects in our Experiment 1 were certainly aware that four different colors were presented in each display and would have noticed if we ever presented additional colors. Subjects, therefore, had access to the identity of the different colors presented even though they did not have information about the activity level associated with each color. Thus, our summary model is likely to be an oversimplification of the actual representation used for decision-making. This point is further underscored by the fact that when predicting the accuracy in the 2-alternative condition, the summary model showed a slight but systematic overprediction in Experiment 1 but underprediction in Experiment 2 (though it was better calibrated in Experiment 4).

Thus, we do not claim that rich information about the visual scene cannot be quickly and efficiently extracted (it can). What our results do suggest, however, is that decision-making circuits do not create a copy of the detailed sensory representation that can be used after the disappearance of the stimulus. This conclusion is reminiscent of the way deep convolutional neural networks (CNNs) operate: the decisions of these networks are based on compressed representations in the later layers rather than the detailed representations in the early layers^40,41^. In other words, even though CNNs extract complex representations in their later layers, the networks do not perform decision making based directly on the more “sensory-like” representations present in their early layers.

In conclusion, we found evidence from one exploratory (Experiment 1) and two preregistered (Experiments 2 and 3) studies that deliberate decision making for discrete stimulus categories is performed based on a summary of, rather than the whole, sensory representation. A final study (Experiment 4) extended these results to stimuli that give rise to continuous multimodal distributions. Our findings demonstrate that flexible computations may not be performed using the sensory activity itself but only a summary form of that activity.

## Methods

### Subjects

A total of 63 subjects participated in the four experiments (32 in Experiment 1, 10 in Experiment 2, 10 in Experiment 3, and 11 in Experiment 4). Each subject participated in only one experiment. All subjects provided informed consent and had normal or corrected-to-normal vision. The study was approved by the Georgia Tech Institutional Review Board.

### Apparatus and experiment environment

The experiments stimuli were presented on a 21.5-inch iMac monitor in a dark room. The distance between the monitor and the subjects was 60 cm. The stimuli were created in MATLAB, using Psychtoolbox 3^42^.

### Experiment 1

The stimulus consisted of 49 circles colored in four different colors – red, blue, green, and white – presented in a ×7 grid on black background. The diameter of each colored circle was .24 degrees and the distance between the centers of two adjacent circles was .6 degrees. The grid was located at the center of the screen. On each trial, one of the four colors was “dominant” – it was featured in 16 different locations – whereas the other three colors were non-dominant and were featured in 11 locations each. The exact locations of each color were pseudo-randomly chosen so that each color was presented the desired number of times.

A trial began with a 500-ms fixation followed by 500-ms stimulus presentation. Subjects then indicated the dominant color in the display and provided a confidence rating without time pressure.

There were three different conditions in the experiment. In the first condition, subjects could choose any of the four colors (4-alternative condition). In the second condition, after the stimulus offset subjects were asked to choose between only two options that were not announced in advance – one was always the correct dominant color and the other was a randomly selected non-dominant color (2-alternative condition). Finally, in the third condition, subjects were told in advance which two colors will be queried at the end of the trial (advance warning condition). For the purposes of the current analyses, we only analyzed the 4- and 2-alternative conditions. The advanced warning condition and the confidence ratings were not analyzed.

Subjects completed six runs, each consisting of three 35-trial blocks (for a total of 630 trials). The three conditions used in the experiment were blocked such that one block in each run consisted entirely of trials from one condition and each run included one block from each condition. Subjects were given 15-second breaks between blocks and untimed breaks between runs. Before the start of the main experiment, subjects completed a training session where they completed 15 trials per condition with trial-to-trial feedback, and another 15 trials per condition without trial-to-trial feedback. No explicit feedback was provided during the main experiment in any of Experiments 1-4 though the presence of second answers in Experiment 3 served as a form of feedback that those specific trials were wrong. We did not hypothesize that the presence or absence of feedback would alter the results in a systematic way and therefore chose to withhold feedback as in our previous experiments^43,44^.

### Experiment 2

Following our exploratory analyses on the data from Experiment 1, we preregistered two additional experiments (Experiment 2 and 3) (<u>osf.io/dr89k/</u>). These experiments were designed to generalize the results from Experiment 1 and to obtain stronger evidence for our model comparison results on the individual subject level. Consequently, we had fewer number of subjects in Experiments 2 and 3 but each subject completed many more trials. We ended up making 3 deviations from the preregistration: (1) we included a different number of locations for non-dominant items (the preregistration wrongly indicated a number that is impossible given the number of categories and total number of characters), (2) we used a dot for the fixation even though the preregistration indicated that we would use a cross-hairline, and (3) we tested additional models (the preregistration only included the population and summary models from Experiment 1).

The stimulus in Experiments 2 and 3 consisted of 49 characters from among 6 possible symbols – ‘?’, ‘#’, ‘$’, ‘%’, ‘+’, and ‘>’ – presented in a ×7 grid. The symbols were chosen to be maximally different from each other. The symbols’ width was .382 degrees on average and height was .66 degrees on average. The distance between two centers of adjacent symbols was 1.1 degrees. The symbols were presented in white on black background. On each trial, one of the six symbols was “dominant” – it was featured in 14 different locations – whereas the other five were non-dominant and were featured in 7 locations each. The exact locations in the ×7 grid where each symbol was displayed were pseudo-randomly chosen so that each symbol was presented the desired number of times.

Each trial began with a 500-ms fixation, followed by a 500-ms stimulus presentation. The stimuli were then masked for 100 ms with a ×7 grid of ellipsoid-shaped images consisting of uniformly distributed noise pixels. Each ellipsoid had width of .54 degrees and height of .95 degrees, ensuring that it entirely covered each symbol. After the offset of the mask, subjects indicated the dominant symbol in the display without time pressure. No confidence ratings were obtained. The experiment had two conditions equivalent to the first two conditions in Experiment 1. In the first condition, subjects had to choose the dominant symbol among all six alternatives (6-alternative condition). In the second condition, subjects had to choose between two alternatives that were not announced in advance: the correct dominant symbol and a randomly selected non-dominant symbol (2-alternative condition).

To obtain clear individual-level results, we collected data from each subject over the course of three different days. On each day, subjects completed 5 runs, each consisting of 4 blocks of 50 trials (for a total of 3,000 trials per subject). The 6- and 2-alternative condition blocks were presented alternately, so that there were two blocks of each condition in a run. Subjects were given 15-second breaks between blocks and untimed breaks between runs. Before the start of the main experiment, subjects were given a short training on each day of the experiment.

### Experiment 3

Experiment 3 used the same stimuli as in Experiment 2. Similar to Experiment 2, we presented a 500-ms fixation, a 500-ms stimulus, a 100-ms mask, and finally a response screen. Experiment 3 consisted of a single condition – subjects always chose the dominant symbol among all six alternatives. However, on 40% of trials in which subjects gave a wrong answer, they were asked to provide a second answer by choosing among the remaining five symbols. Subjects could take as much time as they wanted for both responses. Subjects again completed 3,000 trials over the course of three different days in a manner equivalent to Experiment 2.

### Experiment 4

Experiment 4 employed a modified version of moving dots stimulus adapted from Treue et al.^8^. Three groups of dots moved in three different directions separated by 120°. Unlike many other experiments with moving dots, here all dots moved coherently in one of the three directions. The dots (density: 7.74/degree^2^; speed: 4°/sec) were white and were presented inside a black circle (3° radius) positioned at the center of the screen on gray background. Each dot moved in one of the three directions and was redrawn to a random position if it went outside the black circle. In each trial, a “dominant” direction was randomly selected, and the two “non-dominant” directions were fixed to ±120° from it. The proportion of dots moving in the dominant direction was individually thresholded for each subject before running the main experiment and was always greater than the proportions of dots moving in each non-dominant direction (which were equal to each other).

Each trial began with a white fixation cross presented for one second. The moving dots stimulus was then presented for 500 ms, followed immediately by the response screen which randomly assigned the numbers 1-3 to the three directions of motion (see **Figure 7a**). Subjects’ task was to press the keyboard number corresponding to the dominant motion direction. Similar to Experiments 1 and 2, there were two conditions: subjects chose the dominant direction of motion among all three directions (3-alternative condition) or between the dominant and one randomly chosen non-dominant direction (2-alternative condition). In the 2-alternative condition, the two available options were colored in white and the unavailable option was colored in black.

Each subject completed three sessions of the experiment on different days. Each session started with a short training session. On the first day, subjects completed six blocks of 40 trials with the 3-alternative condition. The proportion of dots moving in the dominant direction was initially set to 60%. After each block, we updated the proportion of dots moving in the dominant direction such that the proportion of dots for the dominant direction increased by 10% if accuracy was lower than 60% or decreased by 10% if accuracy was greater than 80%. Once the task accuracy fell in the 60-80% range, the proportion of dots moving in the dominant direction was adjusted by half of a previous proportion change. After the six blocks, subjects’ performance was reviewed by an experimenter who could further adjust the proportion of dots moving in the dominant direction. Once selected, the proportion of dots moving in the dominant direction was fixed for all sessions. Each session had 5 runs, each consisting of 4 blocks of 50 trials (for a total of 3,000 trials per subject). The 3- and 2-alternative conditions were presented in alternate blocks with the condition presented first counterbalanced between subjects.

### Model development for studies with discrete categories (Experiments 1-3)

We developed and compared two main models of the decision-level representation. According to the “population” model, decision-making circuits have access to the whole sensory representation. On the other hand, according to the “summary” model, decision-making circuits only have the access to a summary of the sensory representation but not to the whole sensory distribution.

In order to compare the population and summary models, we first had to develop a model of the sensory representation. We created this model using the 4- and 6-alternative conditions in Experiments 1 and 2, and the first answer in Experiment 3. The population and summary models were then used to make predictions about the 2-alternative condition in Experiments 1 and 2, and the second answer in Experiment 3. These predictions were made without the use of any extra parameters.

We created a model of the sensory representation for Experiment 1 as follows. We assumed that each of the four types of stimuli (red, blue, green, or white being the dominant color) produced variable across-trial activity corresponding to each of the four colors. We modeled this activity as Gaussian distributions whose mean (*μ*) is a free parameter and variance is set to one. However, in our experiments, the perceptual decisions only depended on the relative values of the activity levels and not on their absolute values. In other words, adding a constant to all four *μ*’s for a given dominant stimulus would result in equivalent decisions. Therefore, without loss of generality, we set the mean for the activity corresponding to each dominant color as 0. This procedure resulted in 12 different free parameters such that for each of the 4 possible dominant colors there were 3 *μ*’s corresponding to each of the non-dominant colors. Finally, we included an additional parameter that models subjects’ lapse rate. Note that the inclusion of lapse rate has a greater influence on percent correct in the 2-alternative compared to the 4-alternative condition because overall performance is higher in the 2-alternative condition. Therefore, introducing a lapse rate favors the population model by leading to predictions of lower performance in the 2-alternative condition (which helps the population model since it consistently predicts higher performance that what was empirically observed).

The sensory representation was modeled in a similar fashion in Experiments 2 and 3. In both cases, the model was created based on subjects choosing between all available options (i.e., the 6-alternative condition in Experiment 2 and the first answer in Experiment 3). The model of the sensory representation in Experiments 2 and 3 thus had 30 free parameters related to the sensory activations (such that for each of the 6 possible dominant symbols there were 5 *μ*’s corresponding to each of the non-dominant symbols) and an additional free parameter for the lapse rate.

We modeled the activations produced by each stimulus type separately to capture potential relationships between different colors or symbols (e.g., some color pairs may be perceptually more similar than others). However, we re-did all analyses using the simplifying assumption that when a color is non-dominant, that color has the same *μ* regardless of which the dominant color is. This assumption allowed us to significantly reduce the number of parameters in our model of the sensory representation. In this alternative model of the sensory representation, the mean activity for each color/symbol was determined only based on whether that color/symbol was dominant or not. Therefore, we included two free parameters for each color/symbol. However, because of the issue described above (adding a constant to all *μ*’s in a given experiment would result in identical decisions), we fixed one of the *μ*’s to 0. This modeling approach reduced the total number of free parameters to eight in Experiment 1 (seven *μ*’s and a lapse rate) and 12 free parameters in Experiments 2 and 3 (11 *μ*’s and a lapse rate). This modeling approach produced virtually the same results (**Supplementary Figure 5**).

Lastly, we considered two different instantiations of the summary model for Experiment 3. In the first instantiation, which we refer to as the Summary & Random Choice model, it is assumed that when the first answer is wrong, then the subject would randomly pick a second answer among the remaining options. In the second instantiation, which we refer to as the Summary & Strategic Choice model, it is assumed that when the first answer is wrong, then the subject would pick the stimulus category of a randomly recalled symbol from the original ×7 grid that is different from the stimulus category chosen with the first answer. According to this model, the subject would pick the second answer correctly 33.3% of the time and each incorrect symbol will be chosen 16.7% of the time (**Supplementary Figure 4**).

### Model development for Experiment 4

Experiment 4 employed moving dots and required subjects to indicate the dominant direction of motion. Therefore, unlike the tasks in Experiments 1-3 that were based on discrete categories of stimuli, the task in Experiment 4 featured a continuous variable (direction of motion, varying from 0° to 360°). However, despite the continuous nature of the stimulus, the three directions of motion could be easily identified implying that the stimulus resulted in a trimodal sensory distribution^8^. This allowed us to use the same modeling approach from Experiments 1-3 by essentially treating the three motion directions as discrete stimuli. We again developed a model of the sensory representation that was fit to the 3-alternative condition. Unlike Experiments 1-3 where the categories of stimuli were fixed, here the dominant direction of motion was chosen randomly (from 0° to 360°) on every trial. Therefore, the model only had parameters for the heights of the non-dominant and dominant directions of motion. Because, just as in the previous experiments, adding a constant to all both parameters would result in identical decisions, the parameters for the non-dominant direction was fixed thus leaving us with a single free parameter. Once the model was fit to the data from the 3-alternative condition, the population and summary models had no free parameters when applied to the data from the 2-alternative condition.

### Model fitting and model comparison

For all four experiments, we fit the models to the data as previously^45–48^ using a maximum likelihood estimation approach. The models were fit to the full distribution of probabilities of each response type contingent on each stimulus type:

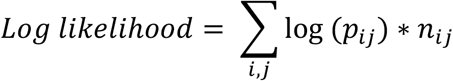

where *p_ij_* is the model’s predicted probability of giving a response *i* when stimulus *j* is presented, whereas *n_ij_* is the observed number of trials where a response *i* was given when stimulus *j* was presented. We give formulas for computing *p_ij_* in a simplified model without a lapse rate in the Supplementary Methods. Because the analytical expressions to obtain *p_ij_* are difficult to compute, we derived the model behavior for every set of parameters by numerically simulating 100,000 individual trials with that parameter set. Model fitting was done by finding the maximum-likelihood parameter values using simulated annealing^49^. Fitting was conducted separately for each subject.

Based on the parameters of the model describing the sensory representation, we generated predictions for the 2-alternative condition (Experiments 1, 2, and 4) and the second answer (Experiment 3) for both the population and summary models. These predictions contained no free parameters. To compare the models, we calculated the log-likelihood ratio (log (ℒ)) of each model. We additionally computed Akaike Information Criterion (AIC):

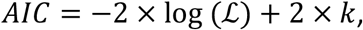

where *k* is the number of parameters of a model. When applied to the 2-alternative condition (Experiments 1, 2, and 4) and the second answer (Experiment 3), the population and summary models had no free parameters. Therefore, AIC was equal to −2 × log (ℒ). We chose to report AIC values instead of the raw log (ℒ) values because of their wider usage and larger familiarity but all conclusions would remain the same if the raw log (ℒ) values are considered. Further, because all of our models had no free parameters, other measures, such as the AIC corrected for small sample sizes (AICc) or the Bayesian Information Criterion (BIC), would result in the exact same pattern of results. Note that lower AIC values correspond to better model fits.

### Data and code

The data from the four experiments, together with all of the analysis codes are freely available online at https://osf.io/d2b9v/files/.

## Acknowledgements

We thank Tristan Hackman, Megan Kelley, and Milan Patel for help with the data collection. This work was supported by the National Institute of Mental Health of the National Institutes of Health under Award Number R56MH119189 to D.R.

## Supplementary material

### Supplementary Methods

#### Mathematically deriving the difference in accuracy between the population and summary models in the 2-alternative condition

In the Results section on Experiment 1, we described the intuition regarding why the population model predicts higher accuracy than the summary model in the 2-alternative condition (**Figure 3**). Here we provide a precise mathematical formulation. Note that the derivations below assume the absence of a lapse rate, which would act to attenuate but not remove the differences between the accuracy levels predicted by the two models.

Let *p_dom=i_* be the probability that, for a particular set of parameters describing the sensory response, the dominant stimulus produces the *i^th^* highest activation. Then, the accuracy in the 4-alternative condition would simply equal *p_dom=i_* as correct trials require that the dominant color produces the highest activation. We can then derive the accuracy in the 2-alternative condition for the population and summary models.

To compute the accuracy in the 2-alternative condition for the population model, we can derive the expected accuracy when the dominant stimulus produces the *i^th^* highest activation for *i* = 1,2,3,4. When *i* = 1, the dominant stimulus produces the highest activation and the subject is always correct. When *i* = 2, the subject is correct when the alternative option happened to be the stimulus with 3^rd^ or 4^th^ highest activation but is wrong when the alternative option happened to be the stimulus with the highest activation. Because the alternative option was chosen randomly, the probability of being correct in this case is 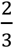. For similar reasons, when *i* = 3, the probability of being correct is 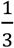 Finally, when *i* = 4, the subject would be wrong regardless which non-dominant stimulus is chosen as the alternative option. Therefore, ACC*_pop−model,2−alt_*, the overall accuracy of the population model in the 2-alternative condition, is:

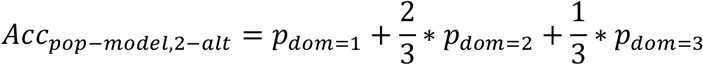

To compute the accuracy in the 2-alternative condition for the summary model, we can again derive the expected accuracy when the dominant stimulus produces the *i^th^* highest activation for *i* = 1,2,3,4. As with the population model, when *i* = 1, the dominant stimulus produces the highest activation and the subject is always correct. However, unlike the population model, *i* = 2,3,4 produce the same probability of being correct. Indeed, in all of these cases, the dominant stimulus does not produce the highest activation and the summary model does not have access to the activations other than the highest activation. From the remaining three stimuli, there is a 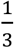 chance that the stimulus with the highest activation was chosen as one of the two options, in which case the subject is always wrong. On the other hand, with 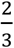 chance another stimulus that did not produce the highest activation is chosen as the alternative option. Because the subject chooses randomly in this case, the probability of being correct is 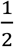. Therefore, the probability of being correct for *i* = 2,3,4 is always 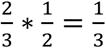 and *ACC_summary−model,2−alt_* the overall accuracy of the summary model in the 2-alternative condition, is:

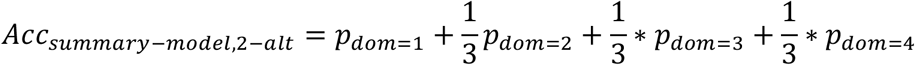

From here we obtain that the difference between the accuracy of the population and summary models in the 2-alternative condition is 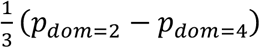. Because the dominant stimulus is at least as likely to produce the second highest than the 4*^th^* highest activation, *p_dom=2_* − *p*_*dom* = 4_ ≥ 0, which means that the population model predicts higher accuracy in the 2-alternative condition compared to the summary model. Similar derivations can be made for Experiments 2-4 as well.

#### Analytical expression for model behavior

For completeness, we provide formulas for *p_ij_*, the predicted probability of giving a response *i* when stimulus *j* is presented. Note that as in the section above, these expressions assume the absence of a lapse rate. When initially fitting the model to the 4-alternative condition in Experiment 1, *p_1j_* equals:

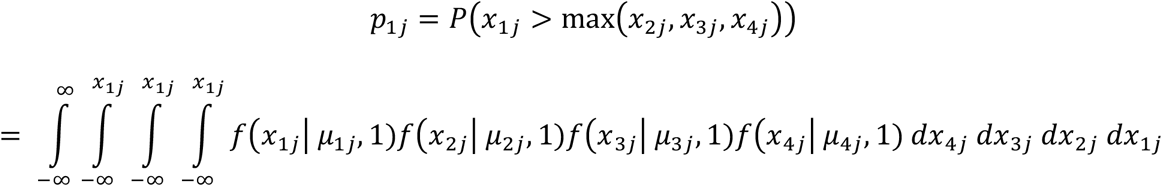

where *μ_ij_* is the mean activity for option *i* when stimulus *j* is presented, *x_ij_* is the activity on a specific trial for option *i* when stimulus *j* is presented, and 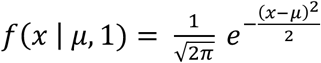 is the Gaussian probability distribution of sensory evidence. The probability **p*_ij_* can be computed in an equivalent fashion when *i* ≠ 1 and when the total number of stimulus categories is different than four (as in Experiments 2-4). Similar formulas can be obtained when fitting the summary and population models to the 2-alternative condition.

#### Model development for all additional models

In addition to the population and summary models, we considered four other models. These models were developed in order to test additional hypotheses about the nature of the representation at the decision stage and the strategies that our subjects could have used. We have not extended this set of four more models even further because any additional models were judged to be too ad hoc and generally provided even worse fits to the data.

The first two of the additional models postulated that decision-making circuits contain information about the sensory representation that is more detailed than the summary model but less detailed than the population model. Specifically, we created models according to which decision-making circuits have access to the highest two or three levels of activation (“2-Highest” and “3-Highest” models, respectively). Just as the summary and population models, these two models were used to predict subjects’ performance in the 2-alternative condition of Experiments 1-2 without any free parameters (the predictions were derived from the same model of the sensory representation used for the summary and population models). Note that the 3-Highest model is functionally equivalent to the population model in the context of Experiment 1 and both the 2- and 3-Highest models are functionally equivalent to the population model in the context of Experiments 3 and 4.

The last two models postulated that subjects attended to just two or three stimulus categories (i.e., colors or symbols) on each trial and made their decisions based on a full probability distribution over the activity levels of the attended categories. We called these the “2-Attention” and “3-Attention” models, respectively. The intuition behind these models is that subjects may not be able to process well the whole set of categories and may therefore choose to focus only on a subset of the categories. The subset was chosen randomly on each trial (otherwise, if subjects always ignored a given stimulus category, that category will never be selected; however, we never observed such behavior in any of our subjects). We first fit these models to the 4-alternative condition (Experiment 1), 6-alternative condition (Experiment 2), and the first answer (Experiment 3) in order to create a model of the sensory representation. The models were then used to predict the 2-alternative condition (Experiments 1 and 2) or the second answer (Experiment 3) without any free parameters. This procedure was equivalent to the procedure used for the summary and population models.

### Supplementary Results

Besides the population and summary models, we constructed and tested four additional models. The “2-Highest” and “3-Highest” models postulated that decision-making circuits have access to the two or three highest activations of the sensory distribution, respectively. These models thus assumed a less severe loss of information compared to the summary model while still postulating that the whole sensory code is not represented in decision-making circuits. On the other hand, the “2-Attention” and “3-Attention” models postulated that subjects choose either two or three stimulus categories to attend to and then make their decisions based on a full probability distribution over the activity levels of the attended categories. We found that neither of these models outperformed the summary model in any of our experiments.

#### Experiment 1

The 2-Highest model (average predicted accuracy = 83.5%) significantly overestimated the observed accuracy level for the 2-alternative condition (average difference = 5.46%, t(31) = 7.49, *p* = 1.94 x 10^−8^) (**Supplementary Figure 1a**). Moreover, the absolute errors in the prediction of the 2-Highest model for the 2-alternative condition (average = 5.86%) is larger compared to the summary model (t(31) = 4.78, *p* = 4.07 x 10^−5^). Model comparison favored the summary model over the 2-Highest model by an average 11.86 AIC points (corresponding to the summary model being 375.41 times more likely for the average subject) and by 379.39 AIC points in the group as a whole (corresponding to the summary model being 2.42 x 10^82^ times more likely in the group) (**Supplementary Figure 1b and c**). Note that within the context of Experiment 1, the 3-Highest model is functionally equivalent to the population model. Indeed, according to the 3-Highest model, the activity level that is not represented is always the lowest; therefore, the 3-Highest model allows one to still order all four activity levels in descending order making it equivalent to the population model.

The 2- and 3-Attention performed even worse. These models could not even be fit to the data in the 4-alternative condition with both models predicting much lower performance (2-Attention model: average difference = −23.8%, t(31) = 22.5, *p* = 9.08 x 10^−21^; 3-Attention model: average difference = −6.73%, t(31) = 10.68, *p* = 6.47 x 10^−12^; **Supplementary Figure 6a**). Nevertheless, we still generated the predictions of these models for the 2-alternative condition and again found that they strongly underpredicted the observed accuracy (2-Attention model: average difference = −21.2%, t(31) = 24.84, *p* = 5 x 10^−22^; 3-Attention model: average difference = −8.36%, t(31) = 11.18, *p* = 2.08 x 10^−12^; **Supplementary Figure 6b,c**). Finally, when compared to the summary model, both models showed much worse fit (2-Attention model: average AIC difference = 1.19 x 10^14^; 3-Attention model: average AIC difference = 8.12 x 10^3^; **Supplementary Figure 6d**).

#### Experiment 2

The 2-Highest model overestimated the observed accuracy in the 2-alternative condition (74.9%; t(9) = 5.65, *p* = 3.12 x 10^−4^) and provided worse fit to the data compared to the summary model (average AIC difference = 18.81 points, total AIC difference = 188.07 points; **Supplementary Figure 2**). The 3-Highest model fared even worse. It overestimated the accuracy in the 2-alternative condition even more severely (76.7%; t(9) = 8.27, *p* = 1.69 x 10^−5^) and provided much worse fit to the data compared to the summary model (average AIC difference = 39.87 points, total AIC difference = 398.65 points; **Supplementary Figure 2**).

Similar to the result in Experiment 1, the 2- and 3-Attention models could not even be fit to the 6-alternative condition with both models predicting much lower performance (2-Attention model: average difference = −21.6%, t(9) = 8.07, *p* = 2.07 x 10^−5^; 3-Attention model: average difference = −11.3%, t(9) = 5.50, *p* = 3.80 x 10^−4^; **Supplementary Figure 6e**). We still generated the predictions of these models for the 2-alternative condition and again found that they strongly underpredicted the observed accuracy (2-Attention model: average difference = − 19.2%, t(9) = 10.25, *p* = 2.91 x 10^−6^; 3-Attention model: average difference = −14.2%, t(9) = 8.35, *p* = 1.57 x 10^−5^; **Supplementary Figure 6f,g**). Finally, when compared to the summary model, both models showed much worse fit (2-Attention model: average AIC difference = 2.71 x 10^61^; 3-Attention model: average AIC difference = 1.30 x 10^37^; **Supplementary Figure 6h**).

#### Experiment 3

The design of Experiment 3 made the 2- and 3-Highest models functionally equivalent to the population model (and thus their predictions were equivalent to that model). Similar to Experiments 1 and 2, the 2- and 3-Attention models did not fit well to the first answer (2-Attention model: average difference = −22%, t(9) = 8.41, *p* = 1.48 x 10^−5^; 3-Attention model: average difference = −11.2%, t(9) = 5.10, *p* = 6.47 x 10^−4^; **Supplementary Figure 6i**). We still generated the predictions of these models for the second answer and again found that they strongly underpredicted the observed accuracy (2-Attention model: average difference = − 23.1%, t(9) = 12.87, *p* = 4.23 x 10^−7^ ; 3-Attention model: average difference = −15.9%, t(9) = 7.46, *p* = 3.84 x 10^−5^; ∼**Supplementary Figure 6j,k**). Finally, when compared to the summary model, both models showed much worse fit (2-Attention model: average AIC difference = 3 x 10^48^; 3-Attention model: average AIC difference = 3.09 x 10^14^; **Supplementary Figure 6l**).

### Supplemenary Figures

**Supplementary Figure 1.**
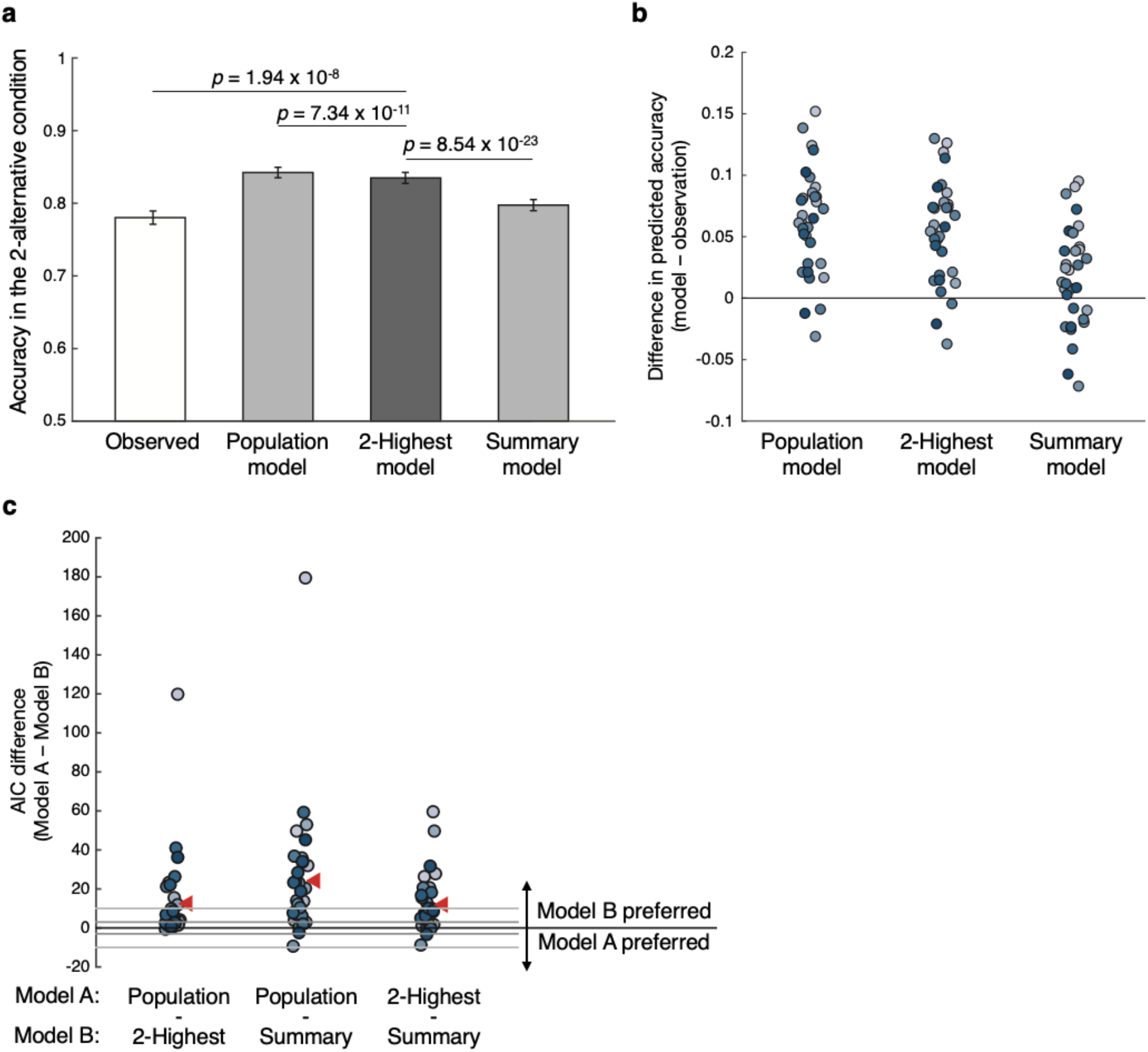
Results for the 2-Highest model in Experiment 1. The results regarding the population and summary models are the same as in Figure 4. (a) Task accuracies of the actual data (white bar; 78% accuracy), the population and summary model (light gray bars) and the 2-Highest model (dark gray bar). The 2-Highest model’s predicted accuracy (83.5%) is in between the accuracy predicted by the population (84.2%) and summary (79.7%) models. (b) Difference in the accuracy for the 2-alterantive condition between the models’ predictions and the observed data. The 2-Highest model’s deviations in predicted accuracy are generally higher than for the summary model. (c) Difference in Akaike Information Criterion (AIC) between the three models. Positive AIC values indicate that the model subtracted (i.e., Model B) provides a better fit to the data. The 2-Highest model provides better fits than the population model, but worse than the summary model in the majority of the subjects.

**Supplementary Figure 2.**
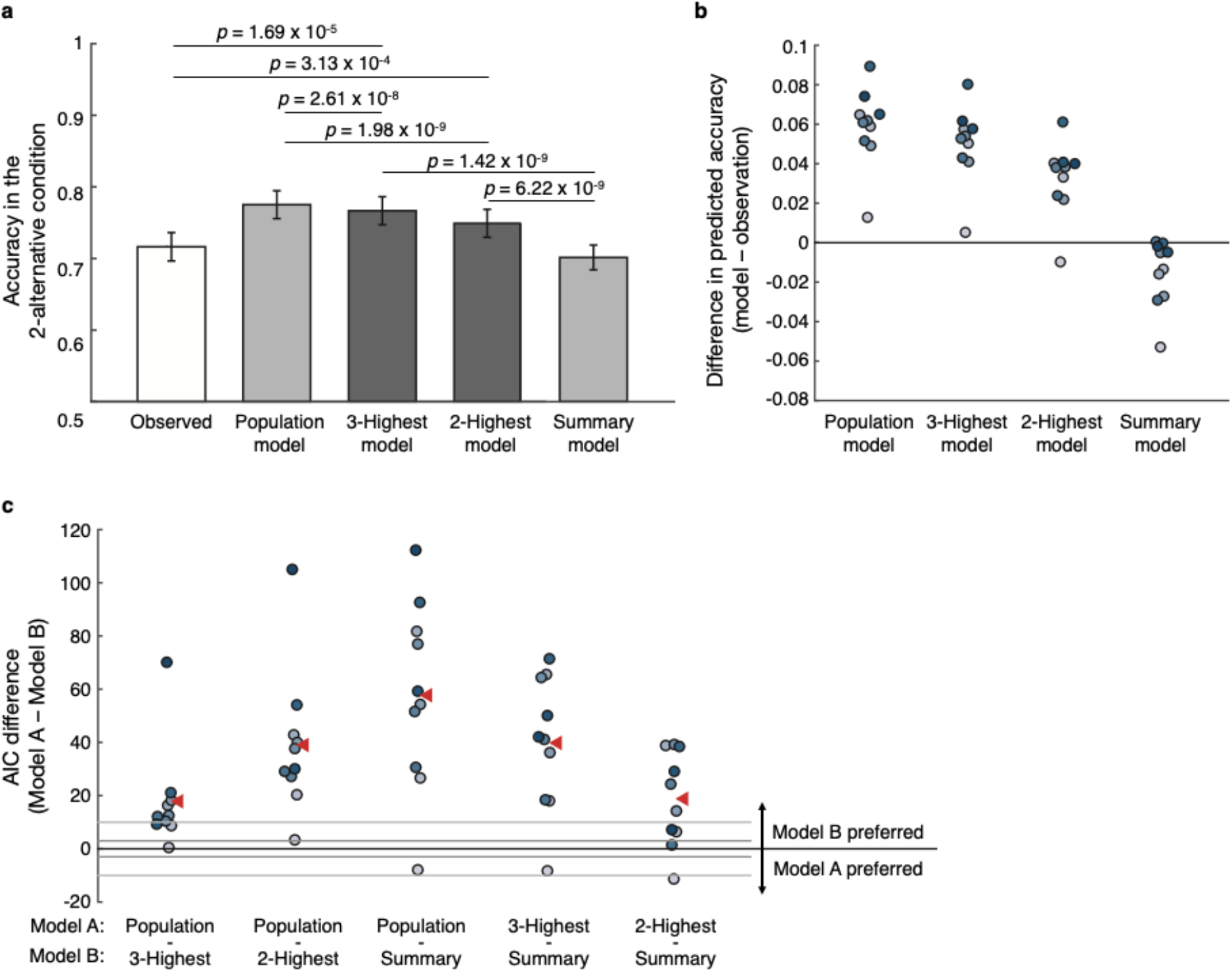
Results for the 2- and the 3-Highest models in Experiment 2. The results regarding the population and summary models are the same as in Figure 5. (a) Task accuracies of the actual data (white; 71.6% accuracy) and the four models. Similar to the result in the Experiment 1, the 2- and the 3-Highest models’ predicted accuracies (74.9% and 76.7% respectively) fell in between the predicted accuracies of the population (77.5%) and the summary (70.1%) models. (b) Difference in task accuracy between the models and the observed data. (c) Model fit comparison between the three models. The summary model has the lowest AIC values, followed by the 2-Highest, the 3-Highest, and the population model.

**Supplementary Figure 3.**
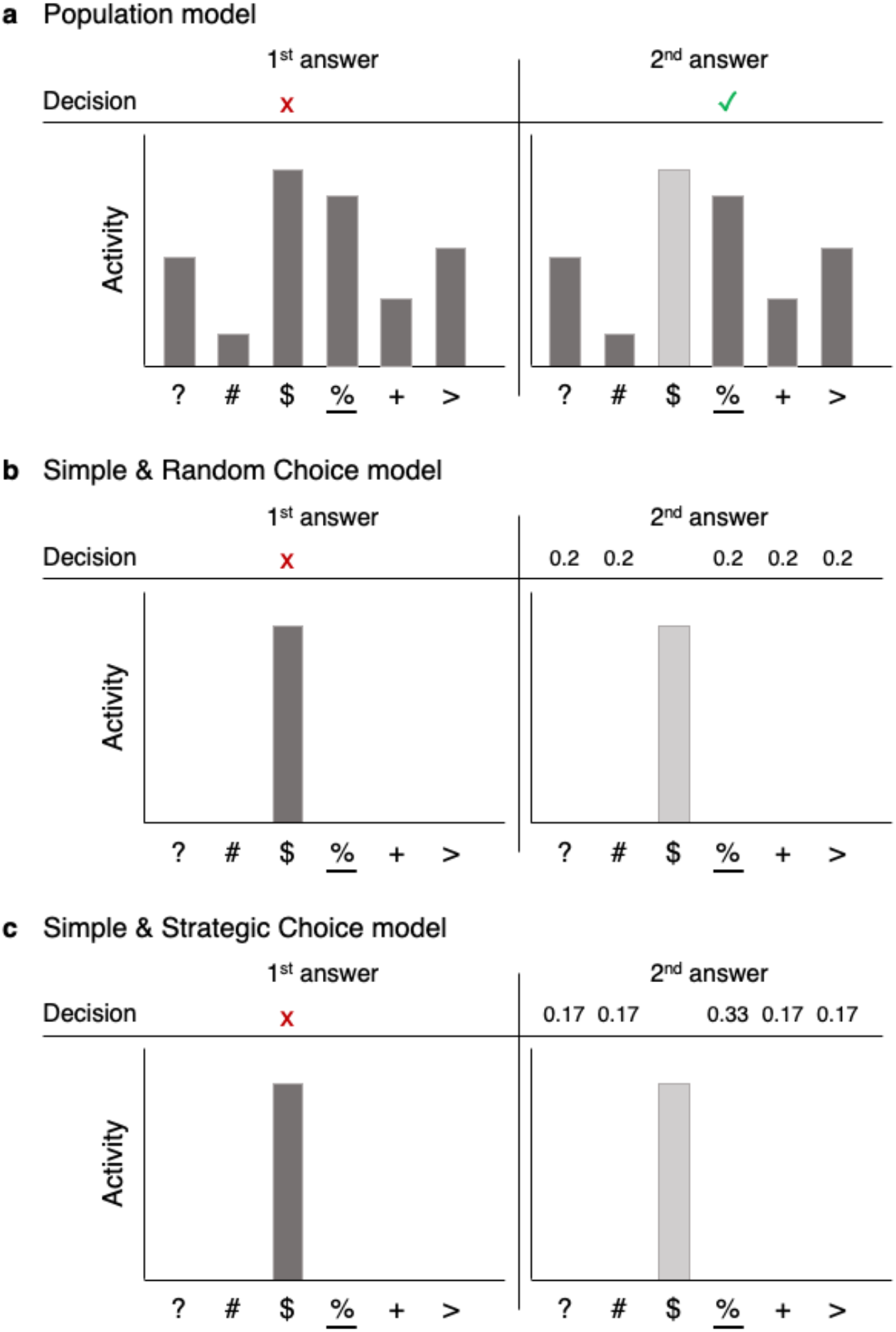
Predictions of the population, the Summary & Random Choice, and Summary & Strategic Choice models for the second answer in Experiment 3. In all examples, the most frequently presented symbol is ‘%’ but the symbol ‘$’ produced the highest activity. The first answer is always the same across all models (left panels) – each model postulates that the highest activity will be chosen first. The predictions diverge for the second answer (right panels; the activity for ‘$’ symbol is represented in a light gray bar to indicate that it cannot be chosen again). (a) According to the population model, decision-making circuits have access to the activity levels associated with all symbols (dark gray bars). Therefore, the population model would imply that the second answer will have a relatively high accuracy since the dominant symbol is likely to have higher activity than the other symbols. (b) According to the summary models, decision-making circuits do not have information about anything but the most highly activated symbol. After an incorrect response, according the Summary & Random Choice model, subjects pick an answer randomly, resulting in 20% accuracy level. (c) According to the Summary & Strategic Choice model, subjects choose the second answer strategically. Specifically, the model postulates that subjects choose the stimulus category of a randomly recalled symbol from the ×7 grid (see Supplementary Figure 4 for details). Therefore, according to the Summary & Strategic Choice model, the accuracy for the second answer will be 33.3%.

**Supplementary Figure 4.**
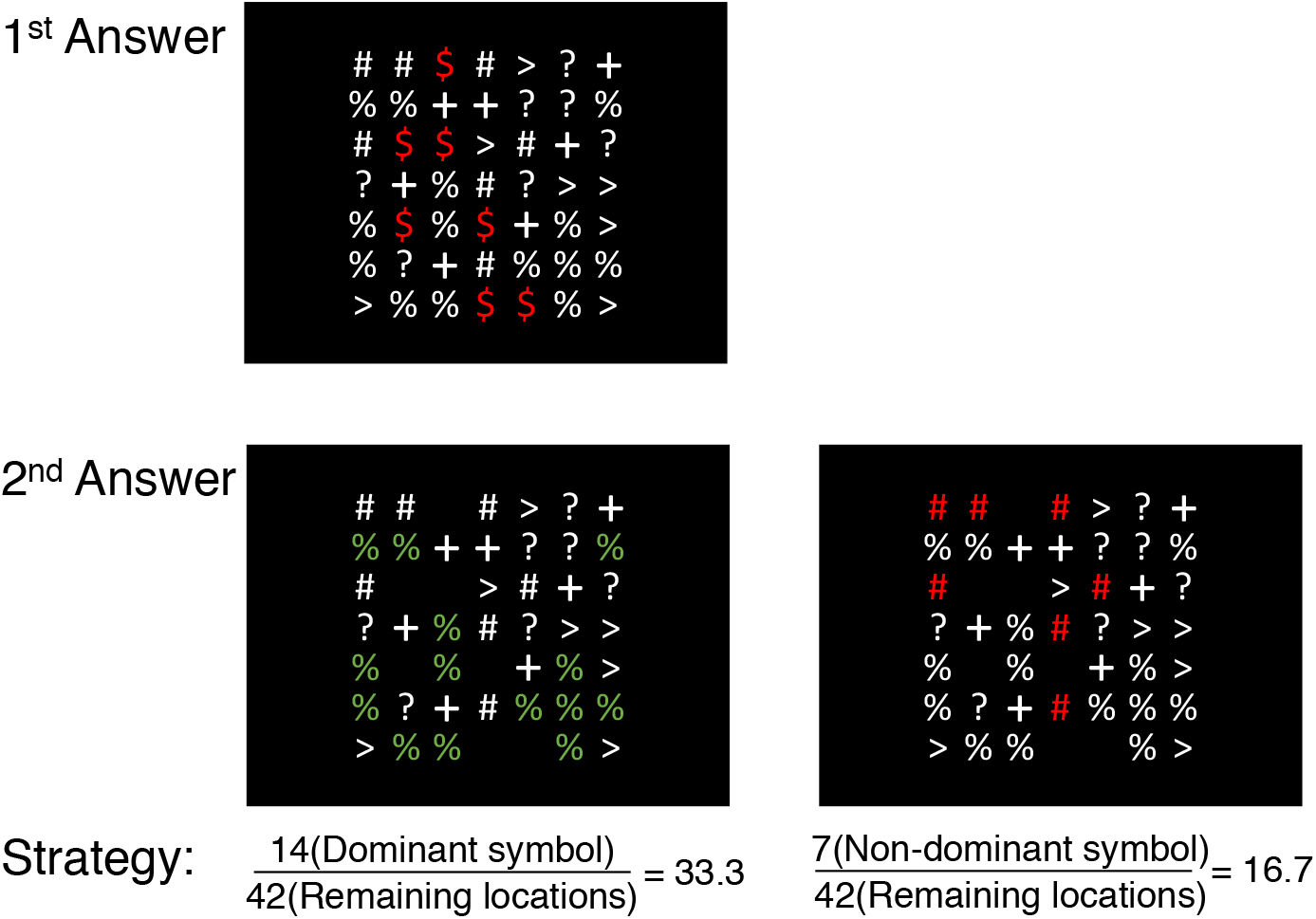
Strategy assumed for the Summary & Strategic Choice model. In the example above, a subject incorrectly chooses the ‘$’ symbol with their first answer (top panel; the correct answer is ‘%’). The model assumes that for their second answer, subjects recollect a single symbol from the original display that was not their first choice and respond with it. Indeed, given that subjects inspected the stimuli for 500 ms, they could easily remember one location with a symbol other than the one they picked for their first answer. With this strategy, the probability that the second answer would be correct is 33.3%, since there were 14 instances of the dominant symbol and 42 locations in the grid (discounting the locations occupied by the symbol chosen with the first answer). Similarly, the probability of picking any specific wrong symbol as the second choice would be 16.7%, since there were 7 instances of that non-dominant symbol and 42 total remaining location in the grid.

**Supplementary Figure 5.**
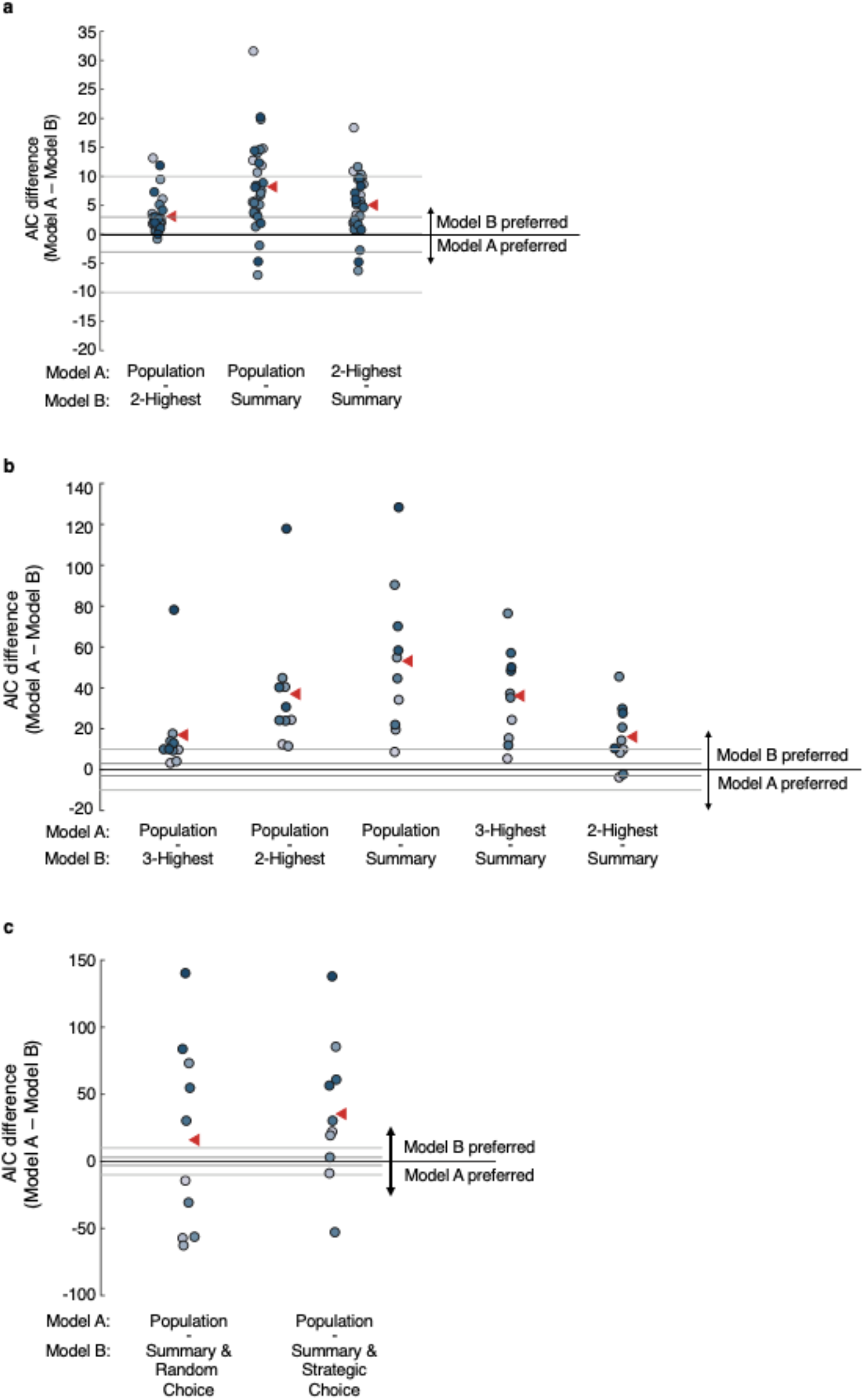
Results of an alternative way of modeling the sensory response. In the analyses reported in the main paper, we modeled the activation levels of each stimulus category (i.e., colors in Experiment 1 and symbols in Experiments 2 and 3) differently depending on the identity of the dominant stimulus. The number of free parameters was thus 13 in Experiment 1 (4 possibilities for the dominant color × 3 free parameters to model the activation for each stimulus category + 1 lapse rate) and 31 in Experiments 2 and 3 (6 possibilities for the dominant symbol × 5 free parameters to model the activation for each stimulus category + 1 lapse rate). We re-analyzed our data using the simplifying assumption of independence between the activations produced by a given stimulus category and the identity of the dominant color. In other words, for example, the color green when non-dominant was assumed to produce the same average activation regardless of whether the dominant color was red, blue, or white. This simplifying assumption decreased the number of free parameters significantly: There were eight free parameters in Experiment 1. The free parameters were used to model the activations of 4 stimulus categories × 2 possible states (dominant/non-dominant) and an additional parameter was used for the lapse rate. However, because adding a constant to all activation parameters retains the relationship between them, one of these parameters was set as zero, bring the total number of free parameters to eight. Similarly, the six symbols in Experiments 2 and 3 resulted in 12 total free parameters. The figure shows model comparison results for this simplified modeling architecture for (a) Experiment 1, (b) Experiment 2 and (c) Experiment 3. In all cases, the summary model is preferred over the population model, typically to the same extent as in the main analyses. Thus, despite the differences between the simplified modeling architecture and the analyses reported in the main experiment, both led to essentially the same results.

**Supplementary Figure 6.**
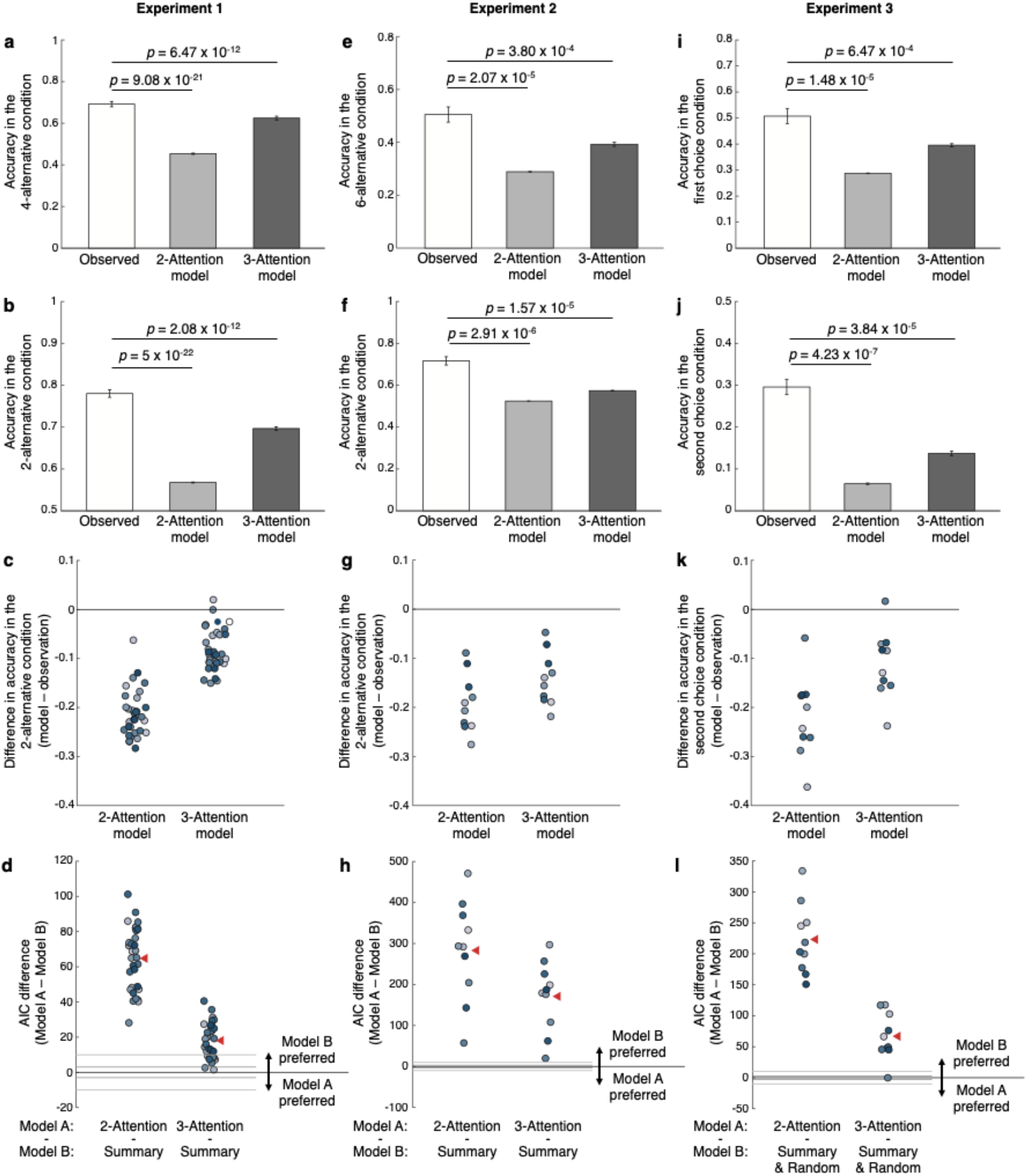
Results of the 2- and 3-Attention models. Both models provided very poor fits to Experiment 1 (panels a-d), Experiment 2 (panels e-h), and Experiment 3 (panels j-l). Specifically, the models could not even fit well the 4-alternative, 6-alternative, and first choice conditions in Experiments 1, 2, and 3, respectively (panels a, e, and i). Not surprisingly, the models also performed poorly for estimating task performance in the 2-alternative (Experiment 1 and 2) and second choice (Experiment 3) conditions (panels b, f, and j). Specifically, the models underestimated task performance in the 2-alternative and second choice conditions (panels c, g, and k). Finally, the both models had substantially higher AIC values compared to the summary model (panels d, h, and l).

